# Combination of 3D and 2D small and wide angle X-ray scattering imaging reveals diminished bone quality in the superior human femoral neck cortex

**DOI:** 10.64898/2026.03.03.709216

**Authors:** Torne Tänzer, Tatiana Kochetkova, Arthur Baroni, Mathieu Simon, Mads Carlsen, Santiago Fernandez Bordin, Manuel Guizar-Sicairos, Philippe Zysset, Marianne Liebi

**Affiliations:** Center for Photon Science, Paul Scherrer Institute (PSI), Villigen, Switzerland; Institute of Materials, École Polytechnique Fédérale de Lausanne (EPFL), Lausanne, Switzerland; ARTORG Center for Biomedical Engineering Research, University of Bern, Switzerland; MAX IV Laboratory, Lund University, Sweden; Institute of Physics, École Polytechnique Fédérale de Lausanne (EPFL), Lausanne, Switzerland

## Abstract

The human femoral neck is particularly vulnerable to fracture, with failure most often initiating in the superior region. While age-related microstructural changes such as cortical thinning and increased porosity are well established, the contribution of material properties at the lamellar and mineralised collagen fibril (MCF) levels remains poorly understood. Here, regional differences in nanostructural properties of cortical bone from 78 femoral necks obtained from 44 donors aged 54-96 are investigated using a combined 2D and 3D X-ray scattering imaging approach. This approach quantifies MCF orientation and structure averaged over multiple lamellae in large fields of view, capturing tissue heterogeneity through the hierarchical scales. We identified misalignment between the scattering signals arising from the MCF bundles — specifically those associated with mineral inclusions in the collagen fibril gap regions, the mineral nanostructure, and the mineral crystal lattice — suggesting the presence of distinct mineral phases within and around the collagen fibers. Despite substantial intra-sample variability, the superior region displays on average more oblique MCF orientations, larger and thicker mineral platelets arranged in a less-ordered structure, greater misalignment between mineral and collagen at the MCF level, and possibly stiffer collagen fibres, with no significant trends observed with donor age or sex. The cumulative effect of these material property differences may contribute to the increased susceptibility of the superior cortex to compressive failure.

## 1 Introduction

Osteoporosis is a major global health problem, affecting more than 1 in 3 women and 1 in 5 men over the age of 50 [1]. The disease markedly increases the risk of fragility fractures, which occur most frequently in the hip and spine and lead to pain, disability, loss of independence, and increased mortality. The economic burden of the disease on the health care systems of Europe was estimated to be more than 56 billion euros in 2019 [2]. Although bone mass is a dominant predictor of fracture risk, optimal management of osteoporosis requires a deeper understanding of bone strength, including the contribution of bone material properties and age-related changes [3].

Bone is a hierarchical material with structural organization on multiple length scales [4], resulting in a remarkable combination of stiffness, strength, and toughness despite being relatively lightweight [5]. Bone is a composite material consisting primarily of type I collagen and a mineral phase of partially carbonated hydroxyapatite (HA). Collagen molecules, staggered with a periodicity of ∼67 nm, are embedded with nanometer-sized HA platelets to form mineralized collagen fibrils (MCFs). The MCFs align into stacked unidirectional arrays with varying orientations, forming lamellae typically 3-7 µm (Figure 1b). The lamellae are concentrically arranged around the Haversian canals, forming osteons—the fundamental structural units of cortical bone [6].

**Figure 1.**
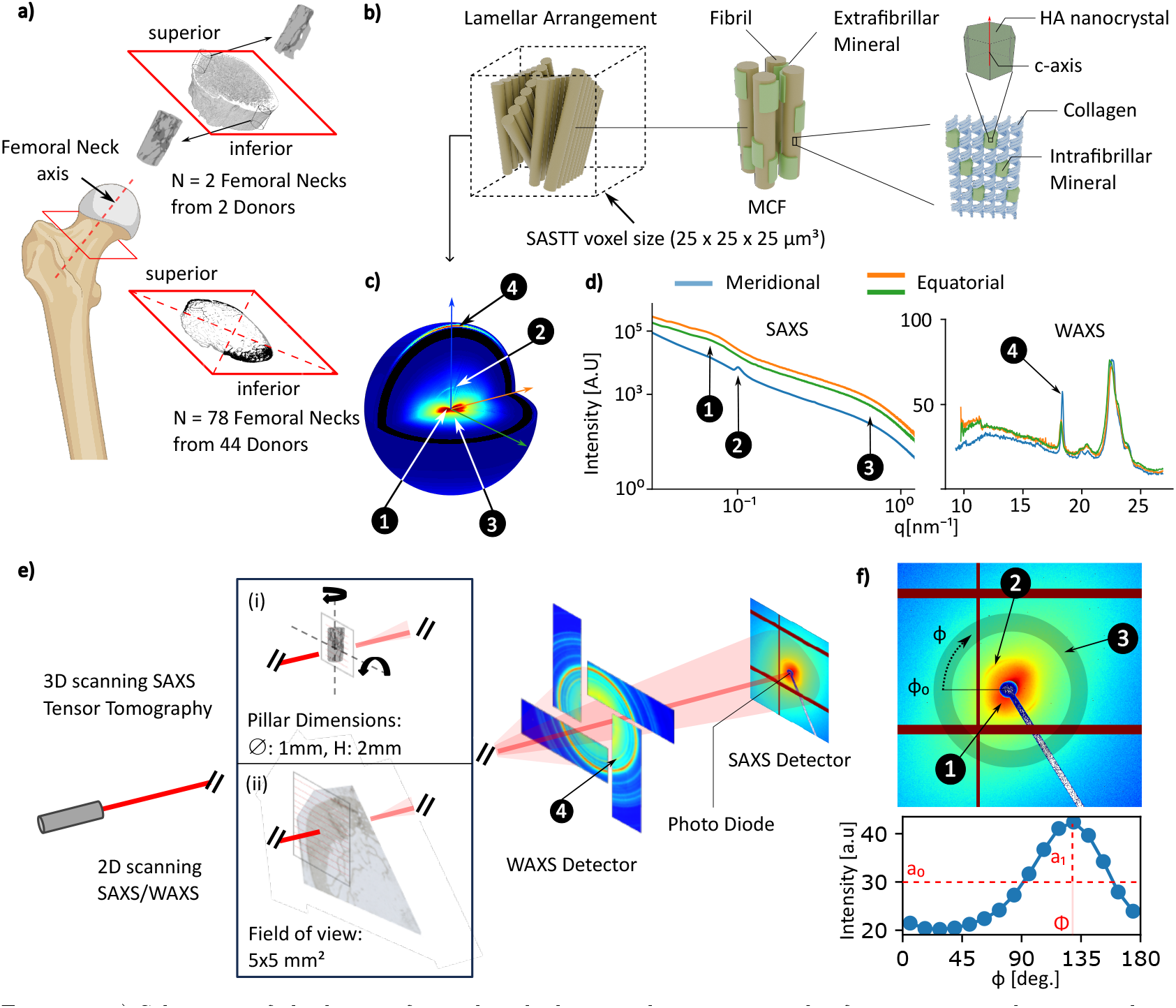
a)Schematic of the human femoral neck showing the superior and inferior anatomical regions, alongwith the locations from which thin sections and micropillars were extracted. b) Schematic representation of bone ultrastructure, showing the hierarchical organization and the relative scale of the imaging voxel (25× 25 ×25 µm^3^). c)Typical 3D RSM, displaying the 4 analyzed scattering signals relating to the MCF organization: collagen equatorial scattering (1), collagen meridional scattering (2), mineral platelet thickness scattering (3), HA (002) diffraction (4). The blue arrow indicates the main MCF direction. d) Scattering intensity along the three orthogonal directions relative to the MCF orientation, indicated by the arrows in (c), plotted across the SAXS and WAXS regimes. e) Schematic representation of SASTT (i) and 2D scanning SAXS/WAXS (ii) experimental setups and location of scattering signal (4) on WAXS (only avaliable in the 2D experiment) detector. f) SAXS detector with location of scattering signals (1-3), and plot of scattering intensity in function of the azimuthal angle *ϕ* in the scattering range (3), indicated by the shaded ring. The mean intensity (*a*_0_), 2D degree of orientation (*a*_1_*/a*_0_) and orientation (Φ) of the scattering signal are indicated.

Consistent with this hierarchical organisation, bone strength is influenced by a wide range of properties across multiple length scales. These include osteonal size and area [7], degree of mineralisation [8], lamellar organisation [9], MCF orientation [10], crystal maturity [11], collagen content [12], enzymatic cross-links [13], non-enzymatic glycation [14], non-collagenous proteins [15], water content [16], as well as diffuse damage and micro-crack density [17, 18]. In recent years, the critical role of material properties at lower length scales has been directly demonstrated through micromechanical testing, including compression [19–21], tensile [22, 23], and bending [24] experiments at the lamellar and MCF levels. Together, these studies highlight the importance of characterising bone structure at these length scales.

The present study focuses on the femoral neck, a key site in hip fracture, which experiences a complex loading environment and distinct loading conditions between the inferior and superior regions (Figure 1a). During gait, the morphology of the femoral neck has led to the hypothesis that the inferior region is predominantly subjected to compressive loading whereas the superior region mainly experiences tensile loading. However, finite element studies that incorporate the effects of muscles and ligaments drawing the femoral head into the acetabulum suggest that most of the femoral neck is in fact loaded in compression [25]. These anatomical locations are also critical during fracture, as most fractures are thought to initiate with compressive failure in the superior region, followed by tensile failure in the inferior region [26]. This underscores the importance of understanding the material properties of femoral neck bone and their relationship to mechanical loading [27]. In particular in the vulnerable superior region, hip fragility has been associated with microstructural material changes such as age related cortical thinning [28], increased cortical porosity [29], and loss of trabecular bone mass and connectivity [30]. However, differences in material properties at the lamellar and MCF levels between anatomical locations and their evolution with age remain incompletely established.

Among the many light, electron, and X-ray based techniques available to investigate the bone material properties at the lamellar scale and below [31,32], small- and wide-angle X-ray scattering (SAXS/WAXS) probes the orientation, organization and dimensions of the MCF (Figure 1b) by statistical average across larger sample areas [33–36]. The scattering signals arising from the MCF are shown in a representative 3D reciprocal space map (RSM) in Figure 1c. The scattering intensity in 3 radial directions oriented with respect to the MCF orientation are shown in Figure 1d. The blue curve represents the scattering intensity in the direction of the MCF orientation, called the meridional scattering. The green and orange curves show the scattering in directions orthogonal to the MCF directions, representing the equatorial scattering. The broad scattering reflection (1) in the equatorial direction is related to the packing and spacing of the MCF bundle (∼100 nm) [37, 38], followed by the distinct meridional collagen peak (2) from the D-spacing of the the periodic mineral inclusions within the gap regions of the collagen fibers ∼ 67 nm [39, 40]. At larger scattering vectors, the scattering signal predominantly oriented in the equatorial direction (3) is attributed to the thickness (*T* -parameter) of the mineral particles (∼2-5 nm) [33]. Moreover, by using a model describing the mineral arrangement as stacks of parallel plates, the SAXS curves can be fitted to estimate the degree of short-range ordering and the typical spacing between successive platelets independently of the amount of mineral scattering [34, 41]. In the wide-angle scattering regime, the reflection corresponding to the ∼0.34 nm lattice spacing of the HA (002) planes (4), which are predominantly oriented colinearly with the MCF, is analyzed [42]. The width and position of the peak allow to determine the length of the mineral platelets and the lattice spacing [35].

The 2D imaging capabilities enabled by scanning SAXS/WAXS [43] have allowed to perform spatial mapping of the aforementioned parameters and have previously been employed to investigate the material properties of the femoral neck. A first study investigated the *T* -parameter in the inferior and superior regions of osteoporotic fracture cases and controls without observing any significant differences, suggesting the possibility of a stiffening process in the organic matrix leading to increased hip fragility [44]. Another study identified hypermineralized regions on the periosteal surface of fracture cases, containing smaller (thickness and length) and less ordered bone minerals in which reduced fracture resistance was demonstrated [45].

The limitations of projection effects and incomplete sampling of reciprocal space in 2D measurements lead to the development of a method combining scanning SAXS/WAXS with computed tomography, called SAXS tensor tomography (SASTT) [46], allowing for the full reconstruction of the 3D reciprocal space in each voxel of 3D samples. Despite the method leading to promising 3D characterization of bone [47, 48], these studies are typically limited by relatively low throughput and small sample volumes, which hinders their ability to capture the biological variability necessary for a comprehensive understanding of the underlying structures.

The purpose of the current study was to combine 3D characterization by SASTT on a limited number of samples with a large-scale 2D statistical analysis to investigate the nanostructural parameters of human cortical bone in the femoral neck, placing particular emphasis on the comparison between the inferior and superior quadrants (Figure 1a) and assessing age-related changes across donors. To this end, 4 pillar samples extracted from 2 femoral necks were characterized by SASTT (Figure 1e(i)) incorporating the most recent algorithmic developments required for the reconstruction of fully anisotropic scattering in lamellar structures [49]. The subsequent analysis of the obtained RSM in each voxel provides a framework for stronger interpretation of 2D SAXS/WAXS measurements on the samples of the same dataset, in which only a slice through the 3D RSM is obtained. Specifically, a model is trained to estimate the out-of-plane angle from the synthetic data created in a transversal cut of the RSMs. In addition, the differences between 3D-averaged parameters and 2D averaged parameters in various cuts of the RSMs can be evaluated to estimate projection effects. In the next step, 156 sections from 78 femoral necks were analyzed using scanning SAXS/WAXS (Figure 1e(ii)) and a robust data analysis pipeline was developed and applied to extract the nanostructural parameters, and the aforementioned model and 3D RSM analysis allowed to extend their interpretation to the 3D structure.

This approach enabled the determination of the dependence of scattering parameters on the underlying MCF distribution and revealed a broad range of structural differences in lamellar and MCF organisation between the inferior and superior regions of the femoral neck. Moreover, this work demonstrates how combining 3D and 2D imaging modalities can help overcome both methodological limitations and the inherent biological heterogeneity of bone tissue.

## 2 Results and Discussion

### 2.1 Exploratory SASTT characterization indicates MCF distribution similarities, orientational and structural differences between anatomical locations

Figure 2a shows the reconstructed orientations of the meridional collagen scattering signal for the 4 samples measured by SASTT grouped into two sample pairs (Pair 1 and 2). Each sample pair consists of the inferior and superior sample extracted from the same femoral neck. Each glyph represents the dominant orientation within a voxel, with it’s orientation and color indicating the angular direction. The pillars were extracted parallel to the femoral neck axis. Consequently desaturated colors according to the colorball indicate longitudinal orientations in the femoral neck, whereas saturated colors correspond to transverse orientations. The meridional collagen scattering corresponds directly to the orientation distribution of the MCF within the voxel. Therefore, for a voxel of 25 ×25 ×25 µm^3^ of lamellar bone, containing roughly 3–8 lamellae, the scattering is distributed within the lamellar planes, mirroring the MCF distribution within those planes (Figure 1b). The scattering intensity reaches a maximum in the dominating orientation of the MCF within the lamellar planes, while the minimum is reached in the direction orthogonal to the lamellar planes (Figure SI1). This anisotropic distribution is quantified by the 3D fractional anisotropy (FA), corresponding to the 3D degree of orientation (Figures SI2).

**Figure 2.**
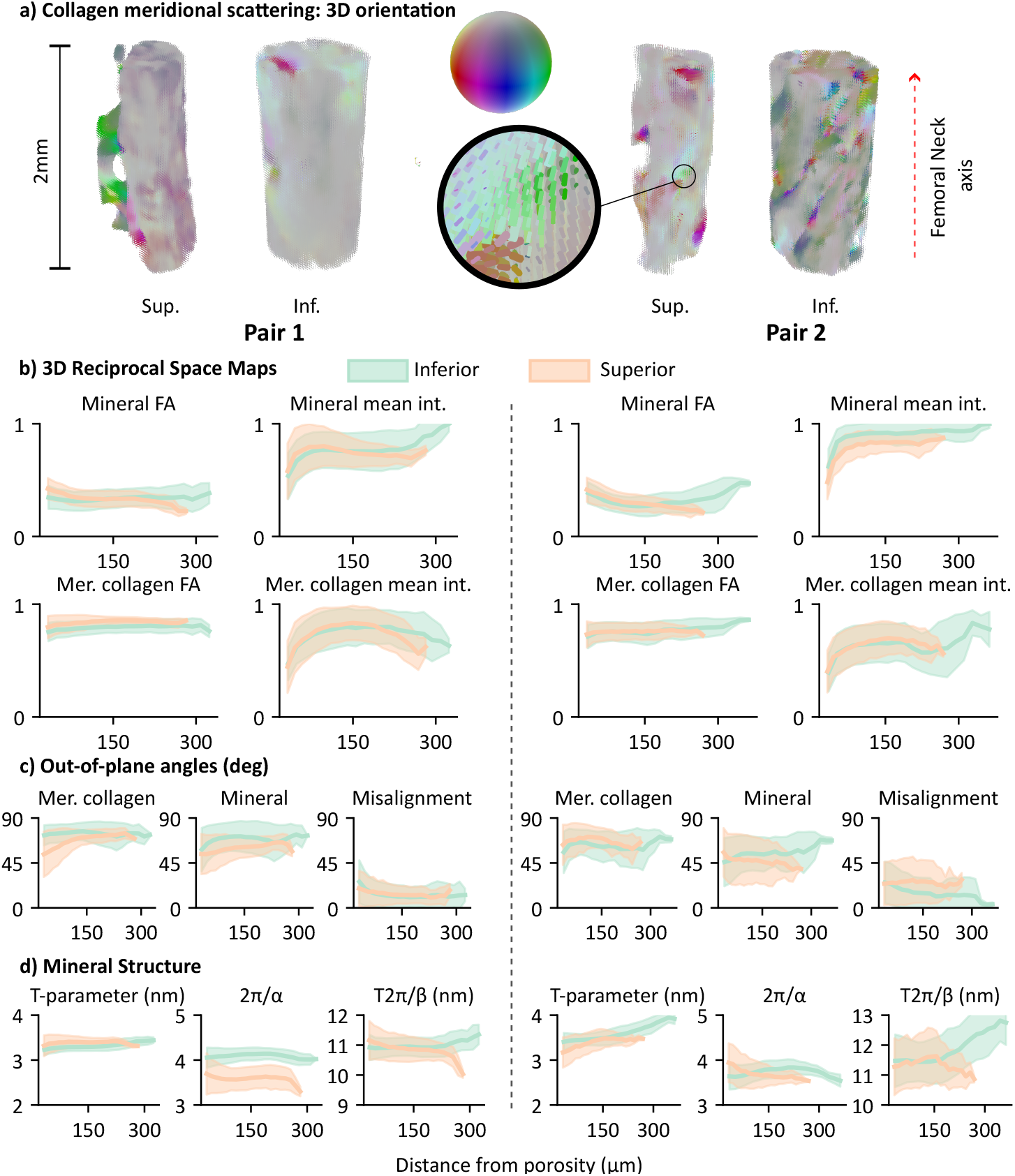
a) 3D glyph plot of meridional collagen scattering orientation in 2 sample pairs. The colorball indicates the 3D orientation: Hue indicates the orientation in the transverse plane, and the saturation indicates the out-of-plane angle. b-d) Mean orientational and structural parameters as function of distance from porosity, seperated by sample pairs (left: Pair 1, right: Pair 2). The intervals correspond to the standard deviation. b) Mean 3D intensity and fractional anisotropy (FA) of the meridional collagen and mineral scattering. c) Out-of-plane angles and angular difference between the meridional collagen and mineral scattering orientations. d) Mineral thickness (*T* -parameter) and arrangement parameters indicating the short-range ordering (2*π/α*) and the average distance between successive platelets (*T* 2*π/β*).

In Pair 1, the superior sample contains more transversely oriented regions and greater spatial variation, whereas the inferior cortex presents a predominantly homogeneous longitudinal alignment (reflected by the desaturated colors according to the colorball). In Pair 2, the superior cortex displays a similar pattern to the superior side in Pair 1, but the inferior cortex shows more heterogeneous orientations, suggesting a more complex underlying microstructure. Overall, the orientations tend to follow microstructural porosities and trabeculae, as previously measured by 3D scanning SAXS [50]. For the quantitative orientational and structural comparison between the anatomical locations, a set of parameters were derived per voxel. The porosity was segmented using the X-ray absorption tomogram and the parameters are presented, per sample pair, as mean values in function of distance from the porosity (Figure 2b-d). With this analysis, the results are decoupled from the microstructural difference between anatomical locations. The median values of the parameters at least 3 voxels (*>*75 µm) distant from porosity are summarized in Table 1. Mean scattering intensities and FA were comparable between anatomical locations in both sample pairs. In all cases, scattering intensities decreased near regions of porosity, likely due to partial volume effects at the bone edges. In particular, the meridional collagen scattering showed no variation between anatomical locations for either pair, and the fractional anisotropy remained constant as a function of distance from the porosity. However, clear changes in orientation of the lamellar pattern were observed both in the transversal (Figure 2a) and out-of-plane directions (Figure 2c). Orientations tended to be slightly less out-of-plane on the superior sides, particularly for the mineral platelet orientation signal. Furthermore, a misalignment between the main orientation of the meridional collagen and the mineral particle scattering was observed, with mean values per sample ranging from 8° to 19° (Table 1). In Pair 2, the misalignment is greater on the superior side, whereas in Pair 1 it is similar between anatomical locations. For the mineral structure, Pair 1 showed similar *T*-parameter and inter-platelet distance values between anatomical locations, but lower 2*π/α* values on the superior side, indicating lower short-range ordering. In Pair 2, the *T*-parameter and inter-platelet distances were higher on the inferior side, while the ordering was similar. Although the *T*-parameter and crystal length have been shown to increase in function of distance from haversian canals [48], there were no systematic changes in function of distance from porosity at this length scale. The collagen D-spacing was equal or lower on the superior side in both pairs (Figure SI3a-b).

**Table 1.**
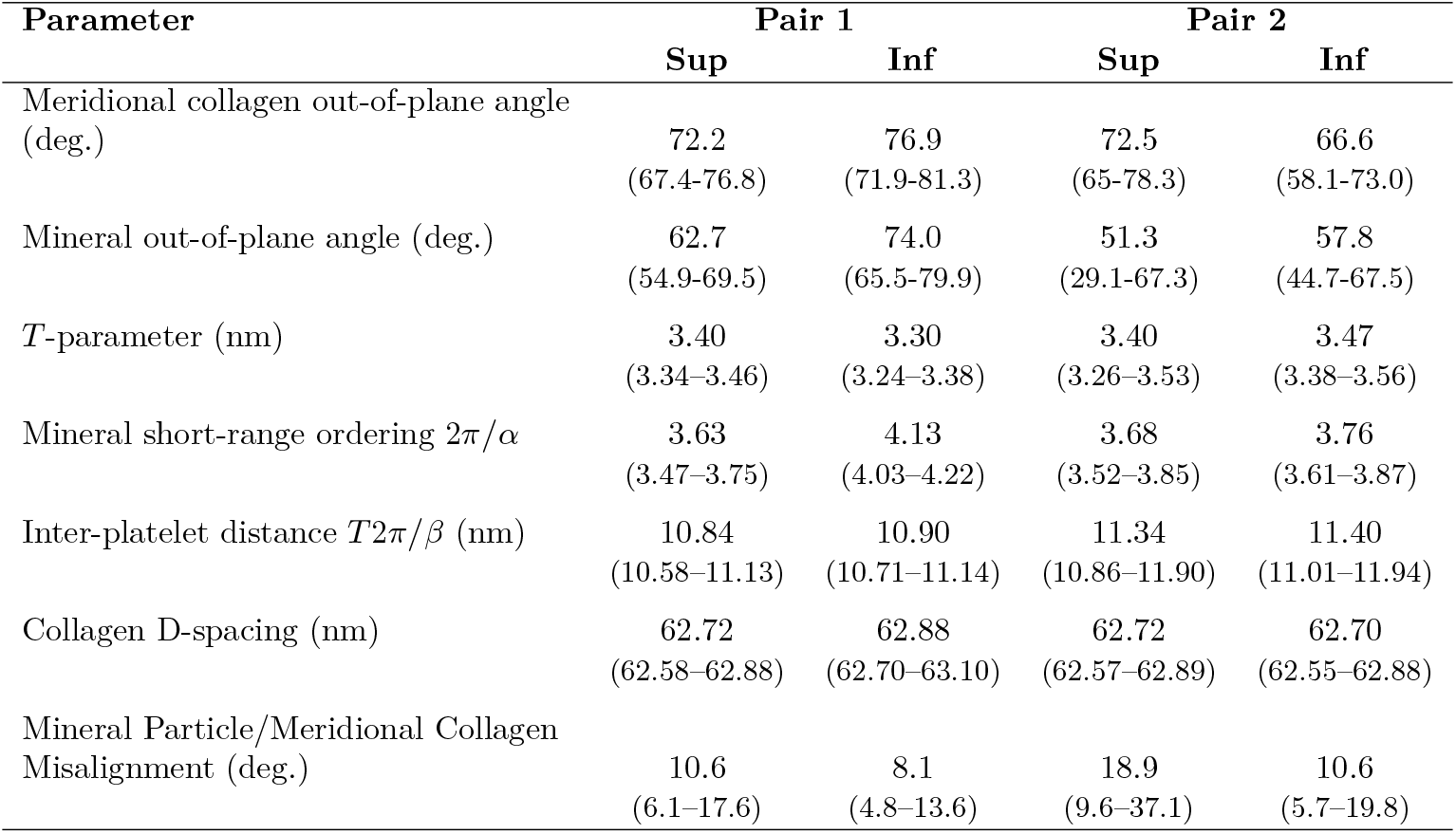
Median values and interquartile ranges (IQR) of the analysed parameters for voxels located at least 75 µm from the porosity in the four SASTT reconstructions.

| Parameter | Pair 1 |  | Pair 2 |  |
| --- | --- | --- | --- | --- |
|  | Sup | Inf | Sup | Inf |
| Meridional collagen out-of-plane angle (deg.) | 72.2<br>(67.4-76.8) | 76.9<br>(71.9-81.3) | 72.5<br>(65-78.3) | 66.6<br>(58.1-73.0) |
| Mineral out-of-plane angle (deg.) | 62.7<br>(54.9-69.5) | 74.0<br>(65.5-79.9) | 51.3<br>(29.1-67.3) | 57.8<br>(44.7-67.5) |
| $T$-parameter (nm) | 3.40<br>(3.34-3.46) | 3.30<br>(3.24-3.38) | 3.40<br>(3.26-3.53) | 3.47<br>(3.38-3.56) |
| Mineral short-range ordering $2\pi/\alpha$ | 3.63<br>(3.47-3.75) | 4.13<br>(4.03-4.22) | 3.68<br>(3.52-3.85) | 3.76<br>(3.61-3.87) |
| Inter-platelet distance $T2\pi/\beta$ (nm) | 10.84<br>(10.58-11.13) | 10.90<br>(10.71-11.14) | 11.34<br>(10.86-11.90) | 11.40<br>(11.01-11.94) |
| Collagen D-spacing (nm) | 62.72<br>(62.58-62.88) | 62.88<br>(62.70-63.10) | 62.72<br>(62.57-62.89) | 62.70<br>(62.55-62.88) |
| Mineral Particle/Meridional Collagen Misalignment (deg.) | 10.6<br>(6.1-17.6) | 8.1<br>(4.8-13.6) | 18.9<br>(9.6-37.1) | 10.6<br>(5.7-19.8) |

The results on the 2 sample pairs highlight the variation between individuals, making it clear that a sufficiently large sample set is needed to obtain statistically meaningful conclusions. To enable optimal analysis of a large-scale 2D scanning SAXS dataset, the influence of the 3D orientation of the scattering signal on the structural parameters evaluated in 2D slices through the 3D RSM was assessed.

### 2.2 Reciprocal Space Map analysis quantifies dependency of scattering parameters on underlying 3D MCF orientation

Figure 3a shows a representative RSM of a 25 ×25× 25 µm^3^ volume of lamellar bone from the SASTT reconstruction (Pair 2 superior). The anisotropy of the scattering signal is illustrated by plotting the scattering intensity in three orthogonal planes (one transverse, two longitudinal), oriented with respect to the femoral neck axis (z-axis) (Figure 3b). The meridional collagen (dashed white circle in transversal cut) is strongest in the longitudinal planes and shows reduced intensity in the transversal plane, as expected with the previously described lamellar structure (Figure SI2). The diffuse fan-shaped scattering from the mineral platelets shows the opposite trend, as expected with the mineral platelets scattering orthogonally to the MCF. The scattering is strongest in the transversal plane and varies significantly between the two longitudinal planes, rather than being axially symmetric, due to the lamellar structure. Note the misalignment between the meridional collagen signal (dashed arrow) and the mineral particle scattering (full arrow).

**Figure 3.**
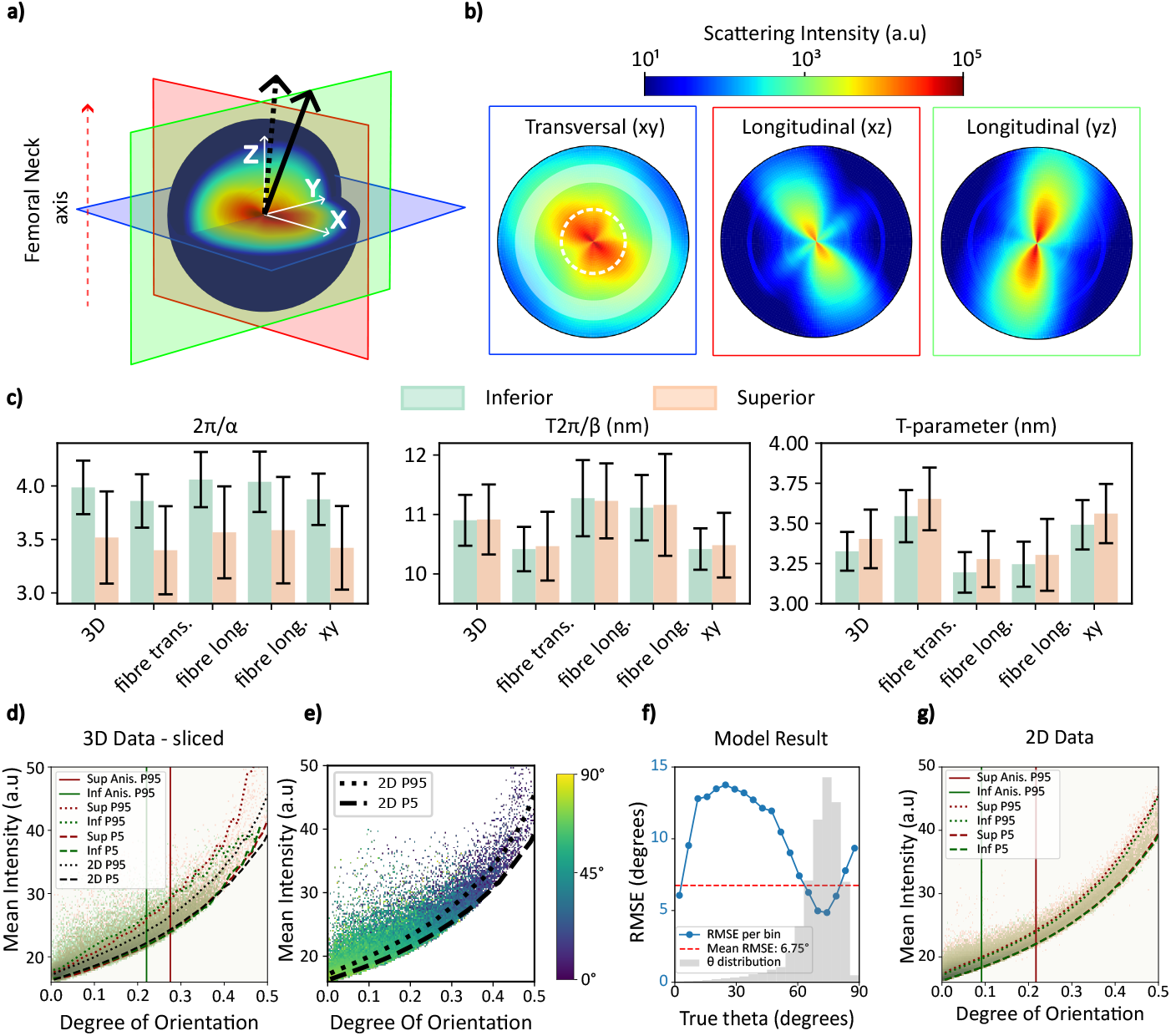
a) Representative 3D RSM of a 25 *×* 25 *×* 25 µm^3^ volume of cortical bone. The three orthogonal planes indicate the transverse (blue, xy) and longitudinal (red, xz and green, yz) planes to the femoral neck axis z. The full and dashed arrows indicate the main orientation of the mineral and meridional collagen (MCF distribution) scattering respectively. b) Representation of the RSM in the 3 orthogonal planes. The diffuse scattering arises from the mineral scattering, and the white dashed line in the transversal cut represents the meridional collagen diffraction peak *q*-range (visible in the longitudinal planes). The shaded ring indicates the mineral *q*-range used to determine the main orientation. c) Comparison between structural parameters, calculated in 3D and in the 3 orthogonal planes. d) Scatter plot of meridional collagen scattering mean intensity vs. degree of orientation of the 3D datasets evaluated in the transversal plane. e)Same as (d), color-coded by the voxel-corresponding out-of-plane angle of the meridional collagen scattering orientation. f) Average RMSE of out-of-plane angle prediction by MLP regression model. g) Scatter plot of meridional collagen scattering mean intensity vs. degree of orientation of the entire 2D dataset.

Under the assumption of large platelets (with platelet length and width not contributing to the scattering in the resolved *q*-range), it has previously been demonstrated that the scattering intensity in any specific direction can be used to analyze the mineral thickness (*T*-parameter) and arrangement parameters of the population of platelets oriented orthogonally to the respective direction [51]. This principle can be extended to different planar cuts through RSMs from SASTT, in which the mean intensity in a plane can be analyzed to obtain the mean parameters of the broader population of platelets orthogonal to the respective plane. For each voxel of the 4 SASTT reconstructions (*∼* 265’000 voxels), the mineral size and arrangement parameters were evaluated separately in the transverse and longitudinal planes with respect to the main MCF orientation in the respective voxel (determined by the meridional collagen scattering orientation) to assess their dependency on fibre orientation. These planes, unique for each voxel, are called the fibre transversal and fibre longitudinal planes. Additionally, the parameters were evaluated in the plane transversal to the femoral neck axis (xy plane in Figure 3a), which is identical for all voxels and corresponds to the plane probed in the 2D experiment. The results are shown in Figure 3c, in comparison to the evaluation on the 3D averaged intensity. In all cases, the 3D values lie between those measured in the separate planes, as expected. In the fibre transversal plane, the *T*-parameter values were on average 8-10% higher than in the fibre longitudinal planes. For the short-range order (2*π/α*) and inter-platelet distance parameters (*T* 2*π/β*), the values are respectively 5-7% and 6-8% lower in the fibre transversal planes. These results indicate that the platelets are thicker and compacter but less ordered in the transversal plane to the main MCF direction, than in the directions along the main MCF orientation. The platelets deviating from the main MCF direction seem to be less thick and compact, but with higher spatial organisation. The differences between the two fibre longitudinal planes are small (∼2%) for all parameters, indicating minimal azimuthal variation around the main MCF direction. Furthermore, for comparison, the parameters are evaluated in the specific directions of strongest and weakest mineral scattering, obtained by Eigenvector analysis of the 3D RSM (Figure SI4). The short-range order and inter-platelet distance were on average up to 18% and 38% higher in the weakest than in the strongest scattering direction, while The *T*-parameter was up to 25% lower. These values display the extent of the 3D directional differences in the mineral size and arrangement at the 25 µm length scale. Lastly, the values measured in the transversal plane to the femoral neck axis (xy plane, Figure 3c) are within 2% of those measured in the fibre transversal plane, reflecting the main MCF orientation in the longitudinal direction of the femoral neck again. The collagen D-spacing value was also evaluated in the different planes, however no relevant differences were noted (maximum 0.3%, Figure SI3c)

### 2.3 Out-of-plane meridional collagen scattering angle can be predicted from transversal plane scattering measurements

It is of great interest to obtain the out-of-plane orientation of the MCF from a 2D transversal cut orthogonal to the femoral neck axis (Figure SI5). This information enables both the determination of the out-of-plane MCF angle and the correction of the orientation effects on the structural parameters demonstrated in the previous section. In the case of an axially symmetric MCF distribution within the voxels, the out-of-plane angle is directly obtainable from the intensity (or 2D degree of orientation) measurement in the transversal plane, if an identical amount of fibres and distribution between voxels is assumed (Figure SI6). However, due to the lamellar structure probed here, the MCF distribution is not axially symmetric within the voxels and the out-of-plane angle is more difficult to access from a measurement in the transversal plane. This is demonstrated by the broad distribution in the plots displaying the out-of-plane angle against the mean intensity and 2D degree of orientation measured in the transversal plane (Figure SI7). However, by using the combined information from the two aforementioned correlations, the out-of-plane angle becomes predictable, up to limitations imposed by the variations of amount and distribution of MCF within the samples.

The relationship between the 2D degree of orientation and mean intensity in the meridional collagen scattering measured in the transversal plane of each voxel of the four SASTT datasets is shown in Figure 3d, color-coded by anatomical location. For a given 2D degree of orientation, the mean intensity exhibits a distribution characterized by a well-defined minimum and a spread toward higher values, as expected from the non–axially symmetric scattering arising from the lamellar structure (Figure SI8). This confirms that the out-of-plane angle dominates the inter-voxel MCF distribution variations in the transversal measurements. The spread to larger 2D degrees of orientations (95th percentile lines in 2D degree of orientation) on the superior side, matches the observed lower out-of-plane angles at this anatomical location (Figure 2c). Figure 3e shows the same data color-coded by the out-of-plane angle of the reconstructed 3D meridional collagen scattering orientation for each voxel. The data is heavily clustered in the bottom-left region of the plot, as highlighted by the 95th percentile lines of the anisotropy distributions, reflecting the dominant longitudinal fibre orientation in the samples (close to 90° out-of-plane angle).

Using the SASTT reconstructions of the four samples, a multilayer perceptron regressor (MLPRegressor, Scikit-learn [52]) with two hidden layers of 64 neurons was trained to predict the out-of-plane angle based on the 2D degree of orientation and mean intensity of the transversal cut of each voxel, as well as the voxel’s anatomical location and distance from porosity. The latter two parameters were included because they also showed correlations with the out-of-plane angle (Figure SI7). The SASTT data used for training the model was restricted to voxels whose transversal degree of orientation and mean intensity fell within the range of the 2D measurements (dashed lines in Figure 3e), comprising approximately 80% of the total dataset (∼205,000 voxels). Model performance is shown in Figure 3f, displaying the mean root mean square error (RMSE) as a function of the true out-of-plane angle. The mean RMSE across all angles is 6.75°, and the model achieves an *R*^2^ value of 0.66. The RMSE correlates inversely with the amount of training data available for each out-of-plane angle, ranging from less than 6° for angles around 60° to 80° to approximately 12° for angles in the 20° to 50° range.

Feature importance was assessed using permutation analysis, with importance defined as the increase in RMSE upon random permutation of each input feature. The analysis revealed that mean intensity contributes most strongly to prediction accuracy, producing an RMSE increase of 10° when permuted, followed by degree of orientation (∼6°) and distance from porosity and anatomical location (*<* 1°). The higher importance of mean intensity is consistent with its stronger correlation with the out-of-plane angle shown in Figure SI6.

The corresponding mean intensity vs. 2D degree of orientation is shown for all points (∼4.7M pixels) of the 2D dataset, color-coded by anatomical location (Figure 3g). The data show the same characteristics in intensity and 2D degree of orientation distributions as in the transversal cuts of the SASTT voxels. The spread to larger 2D degrees of orientations (95th percentile lines in 2D degree of orientation) on the superior side indicates lower out-of-plane angles. However, the spread toward higher intensities is lower than in the SASTT cuts, and the overall 2D degree of orientation values are lower, which might be caused by noise in the tomographic reconstructions. For comparison, the 5th and 95th percentile lines from the 2D datasets are overlaid on the plots for the 3D data (Figures 3d-e).

### 2.4 2D statistical analysis of 78 femoral neck pairs identifies structural deterioration in the superior region

Given the high intrinsic variability of biological tissue, the two femoral neck pairs measured by SASTT were supplemented with 78 additional femoral neck pairs from 44 donors (in some cases, left and right leg) characterized by 2D scanning SAXS/WAXS. For each femoral neck, a 5× 5 mm^2^ region from the inferior and superior quadrants were measured to provide statistical robustness. Figure 4 presents the 2D imaging results of two sample pairs. The difference in cortical thickness is already apparent in the light microscopy images (Figure 4a). The corresponding linear X-ray absorption coefficients are shown in Figure 4b, with values ranging from 12 to 15 cm^*−*1^ showing minimal spatial variation. Figure 4c displays the combined HSV maps (hue: orientation, saturation: 2D degree of orientation, value: mean intensity) of the meridional collagen SAXS signal. Within the bulk of the cortical bone, degree of orientation and intensity values are low, indicating a predominant fibre orientation along the femoral neck axis (orthogonal to the image plane), consistent with the SASTT observations. In trabeculae and around porosities, fibre orientations align with the microstructure, and both degree of orientation and intensity increase, suggesting a higher amount of in-plane fibres. The 2D degree of orientation and mean intensity parameters were used to estimate the MCF out-of-plane angle via the prediction model derived from the SASTT data. The resulting out-of-plane angle map is shown in Figure 4d and exhibits strong correlation with the meridional collagen combined HSV plot as expected. The left (red) sample pair shows comparable nanostructural features between the two anatomical sites, while the right (blue) pair displays a greater proportion of oblique fibres and thicker mineral platelets (Figure 4d). The mean *G*(*x*) function used for fitting the mineral arrangement parameters and the corresponding images are shown in Figure SI9. Again, the left (red) pair has similar mineral arrangement between anatomical locations, while the disorder is greater and inter-platelet distance larger on the superior side of the blue (right) pair.

**Figure 4.**
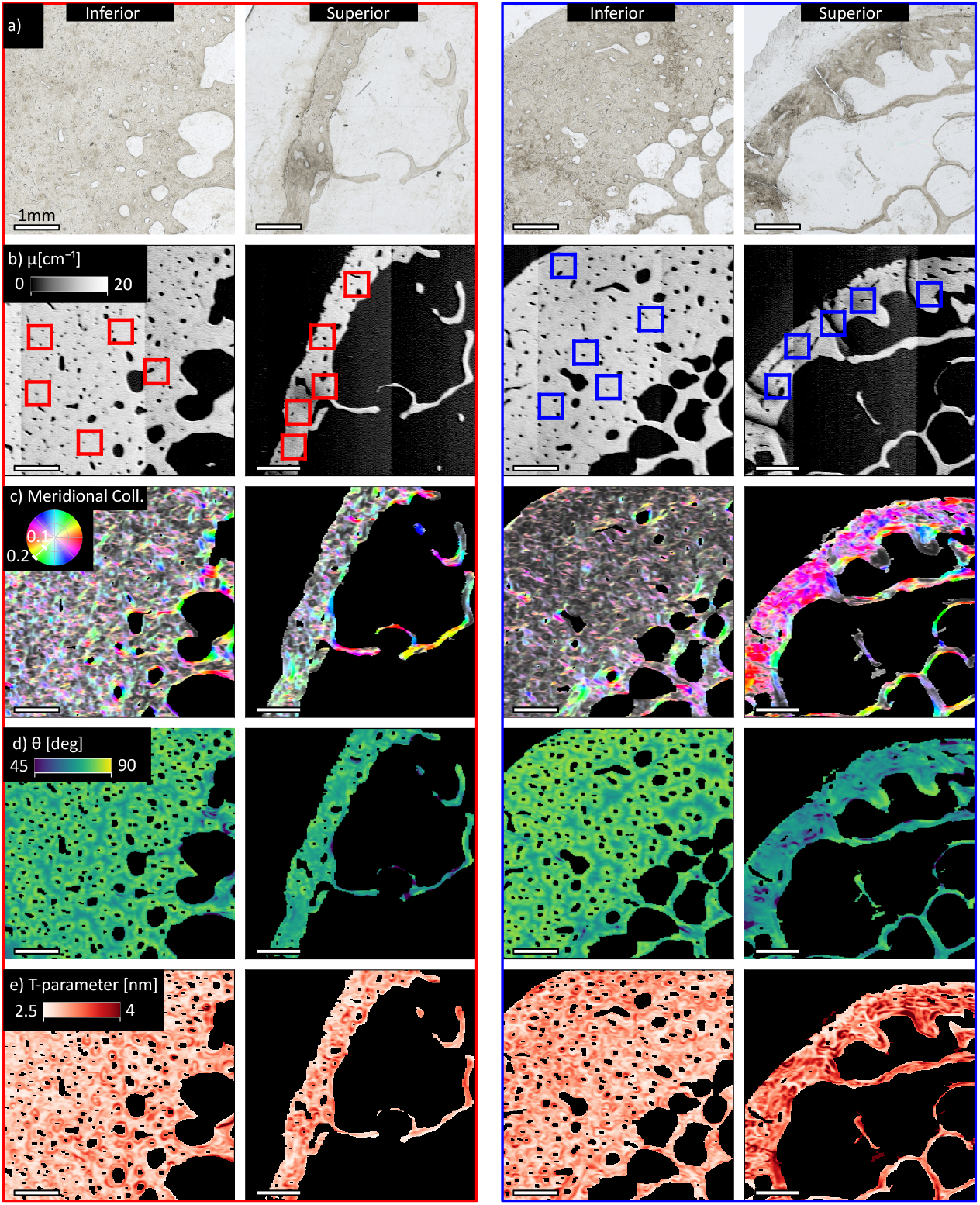
2D imaging of two sample pairs exhibiting minor (red, left) and greater (blue, right) nanostructural differences between anatomical locations. The femoral neck axis is orthogonal to the image plane. a) Light microscopy images. b) Linear absorption coefficient maps. The banding pattern arises from the synchrotron top-up. c) HSV representation of the meridional collagen scattering signal (orientation, 2D degree of orientation, mean intensity). d) Estimated out-of-plane angle of the meridional collagen scattering e) Mineral thickness (*T* - parameter). All scalebars are 1 mm.

For statistical analysis, data from five randomly selected 0.5 × 0.5 mm^2^ ROIs within the segmented cortical bone (red and blue squares in Figure 4b) were included in the linear mixed-effects models, with age, sex, anatomical location, MCF out-of-plane angle and sample thickness as independent variables (see section 5.6). The intraclass correlation coefficients are shown in Tables SI1-2, while the intercept, fixed-effect coefficients and p-values are shown in Table SI3. The two pairs shown in Figure 4 are marked as red and blue points, respectively, in the scatter plots in Figure 5.

**Figure 5.**
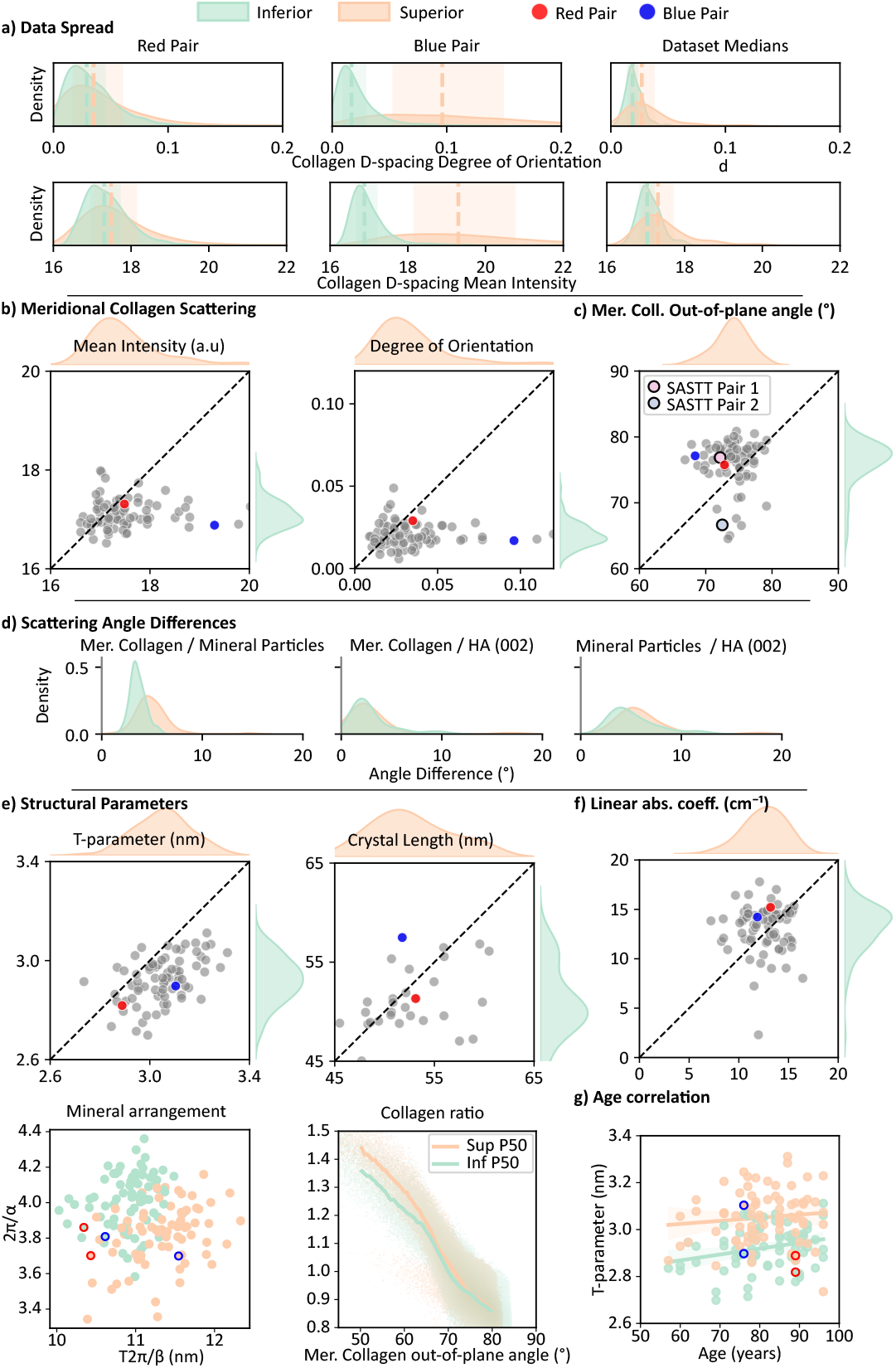
a) Left/middle: Pixel-wise mean intensity and 2D degree of orientation distributions of the meridional collagen scattering for the 2 sample pairs in Figure 4, with median and IQR range for each sample. Right: distribution of sample mean intensity and 2D degree of orientation medians of all 78 femoral neck pairs between anatomical locations. b) Scatter plot of the meridional collagen scattering mean intensity and degree of orientation of all 78 femoral necks, the blue and red points indicate the sample pairs from Figure 4 (x-axis: superior, y-axis: inferior). c) Scatter plot of the estimated median out-of-plane angle of the MCF in the femoral necks. d) Distribution of sample median orientation differences between scattering signals by anatomical location. e) Scatter plots of the median structural parameters (*T*-parameter, crystal length, mineral arrangement, meridional vs. equatorial collagen scattering intensity ratio) of all femoral necks. f) Scatter plot of linear absorption coefficient of all femoral necks. g) Sample median *T*-parameter vs. age of all femoral necks, by anatomical location.

#### Data variability

As expected, pairwise T-tests between measurements from the right and left legs from the same donors showed no difference (*p >* 0.05). In the linear mixed-effects models, variance decomposition for all parameters (Table SI1-2) indicated that donor-level variance (*ICC*_*Donor*_) ranged from 1 % to 20 %, while ROI-level variance (*ICC*_*ROI*_) ranged from 4 % to 19 %. The majority of variability occurred at the pixel level (Residual ×70 % to 95 %), showing that intra-donor variability dominates, with differences between pixels within the same donor and sample greatly exceeding differences between donors. ROI-level variance contributed moderately for some parameters, reflecting heterogeneity across regions within samples. The intra-sample variability of the two example pairs from Figure 4 is displayed in Figure 5a. The pixel-wise distributions of the mean intensity and 2D degree of orientation of the meridional collagen scattering from the highlighted ROI’s are shown in the left (red pair) and middle (blue pair) panels. Despite the large interquartile ranges arising from intrinsic biological variability associated with bone’s hierarchical structure and microstructure-dependent nanostructural organisation (Figure 4), there is a significant difference in the medians between the anatomical locations in the whole dataset (Figure 5a, right).

#### Absorption measurements

The linear absorption measurements are shown in Figure 5f. There is a significant difference (*p <* 0.001) between anatomical locations, with per sample median and interquartile ranges of 12.9 (11.3 - 14.1) cm^*−*1^ on the superior side and 14.0 (11.9 - 14.6) cm^*−*1^ on the inferior side. Previous studies have reported a reduced bone mineral density in the superior quadrant [44, 53, 54]. While the effect of the out-of-plane angle is also statistically significant (*p <* 0.001), its magnitude is negligible compared to that of anatomical location.

#### Mineralized Collagen Fibril Orientations

The 2D degree of orientation and mean intensity of the four analyzed scattering signals (equatorial collagen scattering, meridional collagen scattering, mineral scattering and HA (002) diffraction, see Figure 1), varied significantly with anatomical location. On the superior side, meridional collagen and HA (002) diffraction signals exhibited higher degree of orientation and mean intensity, while mineral and equatorial collagen scattering also showed higher degree of orientation but reduced intensity (Figures 5b and SI10). The relative differences of the per sample medians between anatomical locations are summarized in Table 2. Together, these eight parameters suggest differences in the overall orientation of the MCF between the two anatomical regions, in addition to probable structural changes. As rationalized above, the meridional collagen scattering displayed a strong pixel-by-pixel correlation between mean intensity and 2D degree of orientation (Figure 3f), and was used to estimate the MCF out-of-plane angle. In contrast, the other signals showed much weaker pixel-by-pixel correlations (Figure SI11), suggesting greater variation in the corresponding structural aspects and therefore a weaker direct dependence on the out-of-plane angle. This pattern is also evident in the linear mixed-effects model coefficients: all eight measured parameters (mean intensity and degree of orientation for the four signals) depended significantly (*p <* 0.001) on both anatomical location and MCF out-of-plane angle. For the meridional collagen signal, the mean intensity and degree of orientation showed coefficients roughly an order of magnitude larger with respect to the out-of-plane angle (per 10°), as expected, since these parameters are used to compute the out-of-plane angle. For the other parameters, the effects were distributed between out-of-plane angle and anatomical location. The per sample medians of the MCF out-of-plane angles are shown in Figure 5c. There is a small but significant difference (*p <* 0.001) in the MCF out-of-plane angle between the anatomical locations. The out-of-plane angle is on average 3.1° lower on the superior side, with sample median values of 74.2° and 77.1° on the superior and inferior sides respectively. In the most extreme case, the superior side median was ∼10° lower than the corresponding inferior sample. The intra-sample interquartile range was on average 4.9° and 5.1° in the inferior and superior samples respectively. The orientation of mineralized collagen fibrils (MCFs) in human osteonal bone (femoral midshaft) has previously been quantified using SAXS/WAXS at the lamellar level. These studies employed two-dimensional osteonal sections that were scanned at multiple rotation angles to reconstruct fibril orientation. Wagermaier *et al*. reconstructed the HA (002) orientation and reported angles ranging approximately from 10° to 60°, with a mean around 30° [55]. Using mineral particle scattering, Granke *et al*. similarly observed orientation distributions spanning roughly from 0° to 60° [56]. In both studies, the orientation distributions were asymmetric, consistent with a spiral osteonal architecture rather than an alternating lamellar arrangement [57]. This observation is in agreement with the findings of the present work. To our knowledge, this is the first study to quantitatively report out-of-plane MCF orientations averaged over multiple lamellae. Previous assessments of collagen orientation in the femoral neck have been qualitative, based on polarized light microscopy, and reported brighter, more oblique orientations on the superior side [54]. At the lamellar scale, MCF orientation has been shown to strongly influence compressive mechanical behaviour, with longitudinally oriented MCFs exhibiting higher elastic modulus and yield stress than transversely oriented fibrils [10]. Consequently, the slightly more transverse MCF orientation observed here may contribute to the increased susceptibility to compressive fracture on the superior side.

**Table 2.** Difference (Inf - Sup, %) in mean scattering intensity and degree of orientation between anatomical locations for the 4 analyzed signals, calculated from per sample medians.

| Scattering Signal | Mean Intensity | Degree of Orientation |
| --- | --- | --- |
| Meridional Collagen | -2% | -51% |
| Equatorial Collagen | 4% | -30% |
| Mineral Particle | 20% | -56% |
| HA (002) | 0% | -23% |

#### Inter-signal Orientation Differences

Inter-signal orientation differences were measured to evaluate the MCF organization. Only pixels with a meridional collagen scattering degree of orientation of at least 0.06 were considered, with the aim of avoiding pixels in which the orientation could not reliably be assessed. Figures 5d and SI12 show the angular differences between the signals. The medians of the orientation differences per anatomical location are reported in Table 3. The strongest colinearity occurs between the HA (002) diffraction and the meridional collagen scattering with no variation between anatomical locations, confirming the expected c-axis alignment along the collagen fibers. However there is still a small misalignment, which could be related to the ∼5° tilt of collagen molecules within the fibrils, which means that the overlap zones are tilted with respect to the hole zones, where the inter-fibrillar mineral is located [47, 58]. In contrast, mineral particle scattering displays weaker colinearity with both the meridional collagen and HA (002) diffraction signals, and also varies with anatomical location, showing greater misalignment on the superior side. Also, the superior side shows a greater spread than the inferior side in median misalignment values across samples. Previous high-resolution SASTT measurements have reported local misalignment values of up to 30° between the HA (002) diffraction and the SAXS mineral particle scattering within bands of human lamellar bone [47]. These variations occur on length scales (∼µm) not resolved in the present study, while here we report on misalignment values averaged across several lamellae. The largest misalignment was observed between the collagen equatorial signal and the other three signals (Figure SI12). In these cases, the median misalignment values are slightly higher on the inferior side. However, the broad shoulder of this signal is less reliably extracted from the scattering curves and should therefore be interpreted with caution. The 2D misalignment values only measure the misalignment in the probed transverse plane and are therefore lower than the 3D misalignment values measured in the SASTT reconstructions as expected (Table 1). Note that in the SASTT experiment only the meridional collagen and the mineral particle scattering was in the accessible *q*-range, with misalignment ranging from 5° to 20° in 3D and the 2D projected misalignment ranging from 3.5° to 6.7°. The degree of orientation distributions of the 4 analyzed signals are shown in Figure SI13. Both collagen signals display very low degree of orientation (⪅ 0.15), due to the predominantly longitudinal MCF orientation. Both mineral signals show slightly higher degree of orientation (∼0.2− 0.4), indicating a larger amount of oriented mineral platelets and HA crystals in the transversal plane. This could indicate the presence of extrafibrillar mineral being oriented in the osteonal growth direction.

**Table 3.** Signal misalignment medians and interquartile ranges per anatomical location (deg.), calculated from per sample medians.

| Scattering Signals | Sup | Inf |
| --- | --- | --- |
| Meridional Collagen / Mineral Particles | 4.3 (3.7 - 5.1) | 3.3 (3.0 - 3.8) |
| Meridional Collagen / HA (002) | 2.0 (1.5 - 3.7) | 1.8 (1.5 - 3.9) |
| Mineral Particles / HA (002) | 5.1 (4.2 - 6.3) | 4.2 (3.4 - 6.7) |
| Equatorial Collagen / Meridional Collagen | 12.5 (10.6 - 15.1) | 13.6 (11.8 - 15.6) |
| Equatorial Collagen / HA (002) | 13.4 (11.2 - 15.4) | 14.7 (12.5 - 17.6) |
| Equatorial Collagen / Mineral Particles | 10.8 (9.0 - 12.9) | 12.4 (10.5 - 13.6) |

#### Structural Parameters

The mineral size parameters are increased on the superior side, on average by 4.1% and 5.1% for the *T*-parameter and crystal length respectively (Figure 5e). The mineral short-range order parameter 2*π/α* shows a 4.5% decrease on the superior side, indicating greater disorder, whilst the inter-platelet distance parameter *T* 2*π/β* shows a 3.7% increase on the superior side. These results are represented in the combined mineral arrangement plot color-coded by anatomical location (Figure 5e). The D-spacing of both collagen signals and the HA (002) scattering also showed a small difference (*<* 0.3%) (Figure SI14).

The meridional collagen D-spacing values measured in this study (61-63 nm) are lower than the commonly reported range of 65-67 nm [59], possibly due to sample fixation and preparation effects. As all samples were prepared identically, valid comparisons can nevertheless be made between anatomical locations and donors. The median values and interquartile ranges of the per sample medians of all parameters per anatomical location are provided in Table 4.

**Table 4.** Medians and interquartile ranges of sample medians, per anatomical location.

| Parameter | Inferior | Superior | % Diff (Inf - Sup) |
| --- | --- | --- | --- |
| MCF out-of-plane angle ($^{\circ}$) | 77.1 (75.4–78.2) | 74.2 (72.4–75.5) | **3.8%** |
| $T$-parameter (nm) | 2.92 (2.80–3.08) | 3.04 (2.89–3.23) | **-4.1%** |
| Crystal Length (nm) | 51.67 (45.56–58.28) | 52.92 (46.82–60.91) | **-5.1%** |
| Mineral disorder $2\pi/\alpha$ | 3.97 (3.79–4.14) | 3.79 (3.61–3.96) | **4.5%** |
| Inter-platelet distance $T2\pi/\beta$ (nm) | 10.95 (10.50–11.45) | 11.36 (10.78–11.97) | **-3.7%** |
| Coll. Mer. D-spacing (nm) | 61.31 (60.89–61.68) | 61.12 (60.50–61.57) | **0.3%** |
| HA (002) D-spacing (nm) | 0.34 (0.34–0.34) | 0.34 (0.34–0.35) | **-0.1%** |
| Coll. Equ. D-spacing (nm) | 86.66 (85.33–88.00) | 86.45 (84.70–88.06) | **0.2%** |
| Coll. ratio (Mer. vs. Equ.) | 0.89 (0.85–0.93) | 0.93 (0.89–0.99) | **-5.3%** |

Since the 2D experiment probes reciprocal space in the transverse plane of the femoral neck independently of MCF orientation, and because the 2D structural parameters were shown above to depend on this orientation, the differences observed between anatomical locations must be interpreted with caution. Using the linear mixed-effects models, the corrected difference was derived from the model-predicted values for the inferior and superior locations. These predictions were evaluated at the median values of the remaining independent variables (age, sex, sample thickness, and MCF out-of-plane angle) within each anatomical location. The corrected differences and the fixed-effect coefficient of the out-of-plane angle *β*_4_ are shown in Table 5. Since the calculated out-of-plane angle differs only marginally between locations (∼3°), the corrected differences between anatomical regions change only slightly and remain highly significant (*p <* 0.001), while still showing significant correlation with the out-of-plane angle (Table SI3). For comparison, the out-of-plane fixed-effect coefficient *β*_4_ can be compared to the differences noted in the RSM analysis the SASTT samples, despite the limited range of out-of-plane angles measured in the 2D dataset. For the *T*-parameter, an increase of 0.08 nm*/*10° out-of-plane angle corresponds to a ∼20 % increase in the fibre transversal plane compared to the fibre longitudinal plane, larger than the 8 % to 10 % increase found in the RSM analysis. For the short-range order parameter 2*π/α*, the decrease of 0.02/10° fits very well with the 5 % to 7 % decrease in the fibre transversal plane obtained in the RSM analysis. Finally, the inter-platelet distance parameter *T* 2*π/β* decreases by 1.8 nm*/*10°, leading to ~ 14 % decrease in the fibre transversal plane, slightly larger than the 6 % to 8 % decrease obtained in the RSM analysis. For the D-spacing parameters, differences between anatomical locations and out-of-plane angles remain statistically significant (*p <* 0.001), but their magnitude is minimal (*<* 2 %).

**Table 5.** Percentage difference between Superior and Inferior predicted values from the mixed model and the effect of a 10° change in out-of-plane angle on each parameter.

| Parameter | % Model Diff (Inf - Sup) | $\beta_4$ per $10^{\circ}$ |
| --- | --- | --- |
| $T$-parameter (nm) | **-4.0%** | $7.83 \times 10^{-2}$ |
| Crystal Length (nm) | **-5.1%** | $-3.80 \times 10^0$ |
| Mineral disorder $2\pi/\alpha$ | **4.5%** | $-2.06 \times 10^{-2}$ |
| Inter-platelet distance $T2\pi/\beta$ (nm) | **-3.6%** | $-1.79 \times 10^{-1}$ |
| Coll. Mer. D-spacing (nm) | **0.9%** | $4.41 \times 10^{-1}$ |
| HA (002) D-spacing (nm) | **-0.1%** | $3.00 \times 10^{-4}$ |
| Coll. Equ. D-spacing (nm) | **1.9%** | $1.44 \times 10^0$ |
| Coll. ratio (Mer. vs. Equ.) | **-7.5%** | $-9.72 \times 10^{-2}$ |

The ratio of the meridional and equatorial collagen scattering intensities has previously been linked to collagen cross-linking and fibre-level stiffness [37] and is thus of interest to study. However, this intensity ratio is highly affected by the out-of-plane orientation, which needs to be taken into account for comparing the two anatomical locations. Figure 5e shows this ratio in function of the calculated out-of-plane angle of the MCF for each pixel. As expected, the ratio shows a strong dependence on the out-of-plane angle. However, the median (50th percentile) trends consistently indicate higher values on the superior side. In the linear mixed-effects models, the out-of-plane angle coefficient *β*_4_ corresponds to a decrease of ∼0.1*/*10°, while the superior location contributes an increase of 0.04. These findings therefore suggest a potential stiffening of collagen fibres in the superior region.

#### Age and Sex

In the linear mixed-effect model, none of the investigated parameters showed significant correlation (*p >* 0.05) with age (Figure SI15). We note that although not statistically significant, a small increase in the *T*-parameter with age is observed (superior: 0.013 nm*/*10y; inferior: 0.025 nm*/*10y)(Figure 5g). A previous study reported a logarithmic increase in the *T*-parameter in human cortical vertebral bone between 15 weeks and 97 years [60]. This likely explains why only a small increase was observed in the present cohort, which includes individuals older than 55 years and therefore lacks femoral neck samples from young adults. No differences in X-ray absorption were detected, indicating no measurable change in bone mineral density. The ratio of the meridional and equatorial collagen scattering intensities was previously reported to increase with age [37], but no correlation with age was observed for this ratio in our data. The correlation of all parameters with age are shown in Figure SI15. No significant correlations with sex were noted (Figure SI16).

### 2.5 Nanostructural differences reflect microstructural variations between anatomical locations

The 2D imaging enables spatially resolved mapping of the measured parameters, providing insight beyond the sample-averaged values used in the statistical analysis of the full dataset. The previously reported microstructural differences were observed between the anatomical regions, and appear to be directly related to the nanostructural deterioration. In general, the inferior cortices exhibit a more intact organization, with well-defined circumferential lamellae on the periosteal surface, a well-defined osteonal structure, and a sharp transition to trabecular bone (Figure 6a-c). The meridional collagen scattering and subsequent out-of-plane angle estimation show circumferential lamellae with a more oblique orientation (65° to 70°), while the osteonal bone shows a larger out-of-plane angle (∼80°).

**Figure 6.**
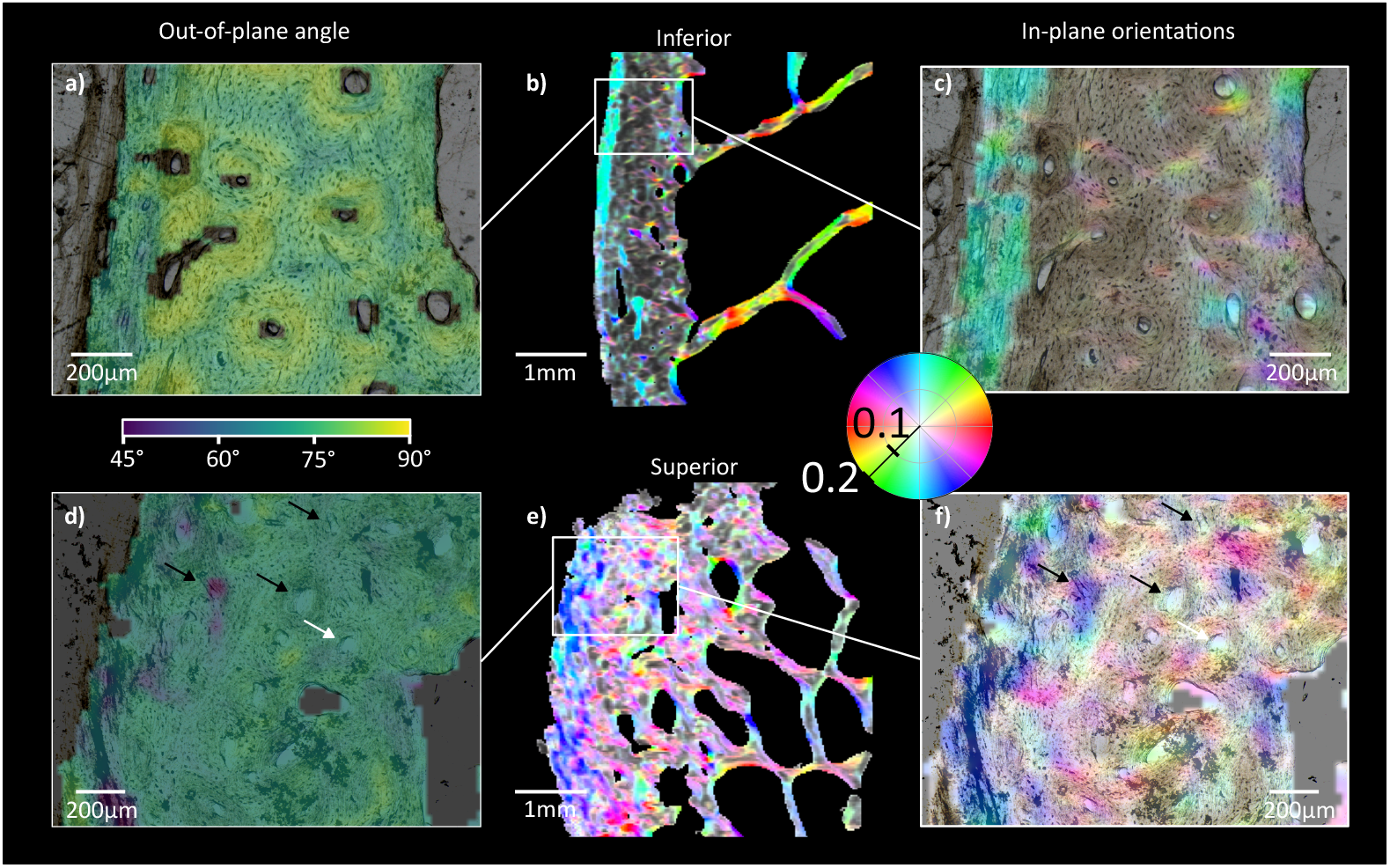
Comparison of in-plane and out-of plane orientations of the meridional collagen scattering of samples overlayed with the corresponding light microscopy images from the inferior (a-c) and superior (d-f) quadrants. (a) The out-of-plane orientations in the inferior quadrant show longitudinally oriented osteons with interstitial bone oriented more obliquely. (b-c) The inferior side presents a clear microstructure, with periosteal circumferential lamellae oriented along the bone surface, a organized cortex of longitudinally oriented osteons showing little degree of orientation (desaturated colors according to colorwheel) and a clear transition to trabecular bone. (d) The out-of-plane orientations on the superior side bone does not present longitudinally oriented bone like on the inferior side, this is due to misaligned osteons (black arrows) but also straight osteons (white arrow) displaying more oblique lamellae orientation. (e-f) The superior microstructure lacks periosteal lamellae, present a disorganized osteonal structure and endosteal resorption.

In contrast, the superior cortices contain extended regions of disordered lamellar tissue, with less pronounced distinctions between osteonal and interstitial tissue, at the microstructural and MCF scales (Figure 6d-f). Osteons are still visible, but they are often misaligned with respect to the longitudinal femoral neck direction, as visible in the microstructure of the light microscope image (black arrows) and in the distinctly lower out-of plane angle of the MCF. However, straight osteons (white arrow) also display more oblique lamellar orientation compared to the inferior osteonal bone. The superior cortices also often display a less distinct transition to trabecular bone, likely associated with endosteal bone loss (Figure 6e and Figure 4 right (blue pair). These microstructural characteristics influence the mechanical behavior of the femoral neck, as periosteal expansion increases cross-sectional area while endosteal resorption reduces it [61].

The present study shows that these microstructural changes are accompanied by changes in the nanostructure, with a rearrangement of orientation of the MCF (Figure 6), but also changes in the shape and arrangement of the mineral particles (*T*-parameter, short-range ordering and average distance between platelets) as can be observed in Figure S17. These structural changes are also noticeable in the blue pair in Figure 4e and Figure S9. Von Kroge *et al*. noted a decrease in the amount of osteocyte lacunae in the superior quadrant [54], which can be related to a reduced capability of mechanosensing and structural adaptation of the bone [62] which could contribute towards a more disorganized structure on the superior side which extends down to the nanoscale.

## 3 Advantages and limitations of this study

The 3D SASTT experiments, which enable reconstruction of the full RSM, allowed us to evaluate how the 2D scattering signals commonly analyzed in bone depend on the underlying 3D orientation of the scatterers. This insight extends the interpretation of 2D scattering data beyond the constraints of the limited reciprocal space probed in 2D experiments, while preserving key advantages such as high-throughput analysis, large fields of view, and the use of thin sections with established preparation protocols. Such sections are widely used in histology, optical and electron microscopy, and mechanical testing, making them well suited for correlative multiscale studies of bone composition, structure, and mechanics. Therefore, the novel combination of 2D and 3D X-ray scattering imaging presented here allows to overcome the key limitations of the methods when used individually. In SASTT reconstructions, regularization is commonly applied to suppress noise. However, its strength must be carefully tuned to avoid overregularization [63]. For structures such as trabecular bone, X-ray absorption contrast and microstructural features can help in assessing the appropriate level of regularization [64]. In compact tissues such as cortical bone, as studied here, this assessment is considerably more challenging. Combining SASTT with 2D measurements on the same samples therefore provides an additional means to evaluate and constrain regularization, enabling more quantitative results.

The prediction of the out-of-plane angle enables to take 2D projection effects into account when analyzing orientation dependent signals, such as the ratio of the meridional and equatorial collagen scattering intensities, but its estimation has several limitations. First, only two femoral neck pairs were analysed using SASTT to estimate orientations in a larger cohort of 78 femoral neck pairs measured in 2D. Second, measurements were performed on transverse sections, whereas MCF are predominantly oriented in the out-of-plane direction in such sections. Longitudinal sections would therefore be more suitable, but are technically challenging to obtain, particularly in the thin superior cortex, and are difficult to define unambiguously, requiring a more precise definition with respect to the femoral neck. As a result, transverse sections are typically analysed. Nevertheless, the present study would benefit from a limited number of longitudinal sections to validate the reconstructed 3D RSMs in the longitudinal plane. Overall, the proposed approach provides an alternative to the direct measurement of out-of-plane angles by sample rotation [55, 65], through the use of complementary 3D measurements.

The *T*-parameter approach used in this study to estimate mineral particle thickness and arrangement depends on the mineral volume fraction, which is assumed to be 0.5. Although X-ray absorption was measured, a quantitative determination of the mineral volume fraction would require calibration using reference samples. Fratzl-Zelman *et al*. quantified calcium weight fractions in the inferior and superior quadrants of the human femoral neck using quantitative backscattered electron imaging and reported a higher mineral content in the inferior region [44], consistent with the X-ray absorption differences observed here. Based on these reported values, a correction factor can be estimated following established approaches in the literature [66]. This analysis indicates that assuming a constant mineral volume fraction equal to 0.5 likely leads to an underestimation of the absolute *T*-parameter values, as well as an under-estimation of the differences in *T*-parameter between anatomical locations. Interpretation of the mineral arrangement in this study is based on the widely used “stack-of-cards” model for platelet organisation, which is well established in the literature [34,41]. Alternative approaches have been proposed that do not require assuming a specific particle shape, such as determining a shape factor and the radius of gyration of the mineral particles as measures of their overall length scale [67, 68]. The shape factor corresponds to the power-law exponent of the scattering curve in the *q*-range 0.18–0.5 nm−^1^ and is presented in Figure SI14, where it shows values very similar to the *α* parameter. In the present work, we adopt the stack-of-cards model, as it enables interpretation of the mineral arrangement in terms of physically meaningful parameters, such as short-range ordering and the average distance between successive platelets.

Finally, the study includes femoral neck samples from donors older than 54 years and therefore does not capture structural differences between anatomical locations at younger ages. In addition, the influence of lifetime physical performance on femoral neck nanostructure would be of considerable interest, as in vivo studies have shown that femoral neck size, bone mineral density, and physical performance together explain a substantial fraction of femoral strength in young men [69]. However, the detailed structural characterisation presented here requires biopsies or post-mortem tissue, for which limited lifestyle information and data protection constraints preclude correlation with physical performance.

## 4 Conclusion

By applying a combination of 3D and 2D SAXS/WAXS imaging to a large cohort of samples (78 sample pairs from 44 donors) from the human femoral neck, various structural differences between the inferior and superior femoral neck cortices were quantified, while accounting for 3D orientation effects. In summary, despite the large intra-sample variability, the on average larger mineral dimensions (*T*-parameter and crystal length), the greater disorder and inter-platelet distances in the mineral arrangement, the lower out-of-plane MCF orientation, the possibly stiffer collagen fibres and the greater misalignment of mineral with respect to collagen fibers, all indicate deteriorated bone quality on the superior side of the femoral neck. While the individual differences are small, the cumulative effect of nanostructural bone property reduction could have an impact on hip fracture susceptibility initiating at the superior side, in addition to the effects of the microstructural and bone mineral density changes. Although the reported parameters reflect nanostructural features at the lamellar and MCF levels, the volumes of interest investigated here span over multiple lamellae. This averaging obscures orientational and structural differences between successive lamellae, but enables parameter mapping over large fields of view, which is essential for characterizing such heterogeneous samples. By covering a large range of scattering angles with two detectors, four relevant scattering features from bone were captured simultaneously. Therefore, in addition to the structural and orientational information from the individual signals, this allowed to identify misalignment between scattering signals at both anatomical locations. Although the true 3D misalignment is underestimated, misalignment between all four signals was measured at both anatomical locations. In particular, the misalignment between mineral particles and collagen indicates a complex organization of the mineralized collagen motif and suggests the presence of distinct mineral phases within and around the collagen fibers.

This work highlights the complexity of structural quantification in biological hierarchical materials such as bone, where advanced combinations of 3D and 2D imaging modalities will continue to offer valuable tools for resolving such challenges in future studies.

## 5 Experimental Section

### 5.1 Sample Preparation

The samples used in this study originate from a set of 78 femurs collected from 44 donors, 45 female and 32 male, all aged between 54-96 years and were obtained with the written consent of the donors by Vienna University hospital. The central part of the femoral necks were imaged by mico-computed tomography and tested mechanically in compression in a previous study [70], whereas the distal sections for the current study (approximately 2 cm in height) were fixed and embedded in poly methyl methacrylate following standard procedures. The four quadrants (inferior, anterior, superior, posterior) were defined by fitting an ellipse and calculating the center of mass of the cross section of the femoral neck, indicating the direction of the inferior side of the femoral neck. The inferior quadrant was defined as the region spanning 60° around the calculated direction, and the superior side as it’s opposite. For the 2D SAXS/WAXS, thin sections were extracted from the inferior and superior quadrants (Figure 1a). The quadrants were cut to approximately 400 µm thick sections using a diamond band saw under constant water irrigation (Exakt, Norderstedt, Reichert-Jung). In the next step, the thick sections were ground to approx. 100 µm thin sections using a Logitech Ltd PM5 precision lapping and polishing system. The final step consisted of manually fine grinding and polishing the samples with a solution of demineralized water and 10% of *Al*_2_*O*_3_ 0.05 µm powder to obtain thin sections in the range of 50-80 µm, as measured with a digital caliper. For the 3D SASTT measurements, 4 micropillars (1 mm diameter, 2 mm height) were milled from the cortical center of the inferior and superior quadrants of two femoral necks using a custom-made lathe system [71] (Figure 1a).

### 5.2 2D SAXS/WAXS

The samples were measured at the ForMAX beamline at the MAX IV laboratory (Lund, Sweden). A double crystal monochromator was used to fix the X-ray energy at 16.13 keV and the beam was focussed to 50× 10 µm^2^ in the sample plane, as measured by knife edge scan. A flight tube was positioned between the sample and the detectors to minimize the air scattering and absorption. The samples were raster scanned with a step size of 25 ×25 µm^2^ over a 5× 5 mm^2^ area in the central part of each quadrant with an exposure time of 0.1 s at each point, and the scattered X-rays were collected on *X-Spectrum Lambda 3M* (WAXS) and *Dectris EIGER2 X 4M* (SAXS) detectors placed 0.134 m and 6.9 m downstream of the samples respectively. The transmitted beam was measured by a photodiode mounted on the beamstop inside the flight tube (Figure 1e). The *q*-range accessed on each detector was determined by calibrating with silver behenate, leading to available SAXS and WAXS *q*-ranges of 0.02 - 1.5 nm^*−*1^ and 12.5 - 26 nm^*−*1^ respectively (Figure 1d).

### 5.3 2D SAXS/WAXS analysis

The 2D scattering patterns collected at each measurement point were normalized by the X-ray transmission and radially integrated into 1D scattering curves across 32 azimuthal segments using PyFAI [72]. This data reduction is particularly valuable for quantifying scattering mean intensity and degree of orientation within specific q-ranges through a Fast Fourier Transform-based approach [43](Figure 1f). The bone tissue was automatically segmented by Otsu’s method [73] and the cortical region was manually defined. For each sample, 5 ROI’s of 0.5 ×0.5 mm^2^ were randomly selected in the cortical region for the characterization of each sample (Figure 4b).

The four distinct scattering features associated with mineralized collagen fibrils are illustrated in Figure 1c-f. In the low *q*-range, the diffraction pattern is dominated by a broad equatorial peak at approximately *q≈* 0.05 nm^*−*1^, corresponding to the lateral packing of collagen fibrils. A sharp meridional reflection at *q ≈* 0.1 nm^*−*1^ reflects the characteristic D-spacing ∼ 67 nm of the periodic mineralization along the collagen fibrils. At *q≈* 0.4 nm^*−*1^, the signal arises from equatorial scattering associated with the thickness of the mineral platelets. In the WAXS regime, a distinct peak at *q≈* 18.4 nm^*−*1^ corresponds to the [002] reflection of hydroxyapatite.

**5.3.1 Collagen signal analysis**

The meridional collagen peak was extracted in the intensity vs. *q* curves of each segment, using a baseline extraction in an extended q-range (q = 0.07 - 0.13 nm^*−*1^). The integrated area, peak amplitude, full-width half maximum and position of the extracted peak were determined by fitting a Lorentzian function. The orientation and degree of orientation of the collagen fibrils was determined using the Fast Fourier Transform method of the extracted signal in the *q*-range of the peak (q = 0.09 - 0.11 nm^*−*1^).

Once the meridional peak was subtracted from the intensity curve, the same peak extraction and fitting approach was used to characterize the equatorial scattering peak in the *q*-range 0.04 - 0.12 nm^*−*1^. The ratio of the integrated areas was calculated.

#### 5.3.2 Mineral Particle Thickness and arrangement analysis

The orientation and degree of orientation of the diffuse scattering originating from the mineral particle thickness was analysed using the Fast Fourier Transform approach in the *q*-range 0.18 - 0.5 nm^*−*1^. The particle thickness and arrangement was obtained by the *T*-parameter method [33, 34, 41]. Assuming a two-phased system with sharp interfaces, the integral parameter *J* = ∫ *q*^2^*I*(*q*)*dq* and the Porod constant *P* = lim_*q*→ ∞_ *q* ^4^*I*(*q*) can be related to the interface surface area to volume ratio *σ* and the phase volume fraction *ϕ* by [74]:

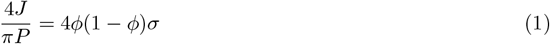

Assuming equal volume fractions (*ϕ* = 0.5) and platelet shaped mineral particles, equation 1 simplifies to:

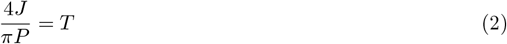

where *T* is the mineral platelet thickness. Since only part of the integral parameter *J* is measured experimentally, the azimuthally averaged curve has to be extrapolated in the low and high-*q* regions missing in the measurement. For the low-*q*, the data is linearly exrtrapolated from *q*_*min*_ to *q* = 0, whereas the extension of the Porod regime to *q*_*max*_ = is used for the high-*q* extrapolation. Following the approach from [33], the spatial arrangement of the mineral crystals is contained in a rescaled dimensionless function of the scattering intensity *G*(*x* = *qT*):

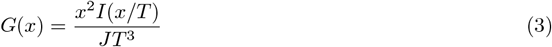

which is directly related to the 1D correlation function *g*(*ζ*) by Fourier Transform:

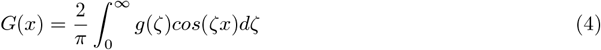

Following the procedure described by Gourrier *et al*. [41], a damped oscillation is considered for the correlation function *g*(*ζ*), resulting in *G*(*x*) taking the form:

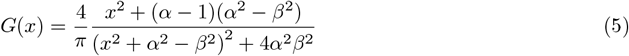

where the parameters 2*π/α* and *T* 2*π/β* describe the short-range ordering and the average distance between successive plates. In practice, these parameters are estimated by fitting the model of *G*(*x*) to the data:

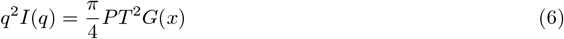

using *α, β* and *T* as fitting parameters.

#### 5.3.3 WAXS analysis

For the WAXS data analysis, only a subset of 31 sample pairs was retained, as sample preparation had affected the WAXS signals in the remaining cases. The hydroxyapatite (002) reflection was extracted from the integrated intensity curves and fitted using a Lorentzian in a *q*-range 18 - 19.4 nm^*−*1^. The peak intensity in each segment was used to extract the orientation of the crystallographic *c*-axis, and the crystal length *D* was calculated using the FWHM of the peak (*B*) and the Scherrer equation:

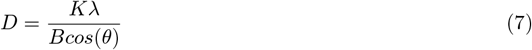

where *λ* is the X-ray wavelength, *θ* the diffraction angle, and *K* a constant usually equal to 1.

### 5.4 SAXS Tensor Tomography Experiment

The micropillars were measured at the cSAXS beamline of the Swiss Light Source (SLS) at the Paul Scherrer Institut (PSI), Switzerland. The X-ray energy was set to 11.2 keV using a Si (1 1 1) double crystal monochromator, and the scattering patterns were recorded on a Pilatus 2M detector placed at a sample to detector distance of 2.17 m. A flight tube, approximately 2 m in length, was placed in between the sample and detector to reduce the air scattering. A 1.5 mm steel beamstop inside the flight tube blocked the directly transmitted beam. The fluorescence signal from the beamstop, proportional to the intensity of the impinging X-rays was measured by a Cyberstar (Oxford Danfysik). This allowed the relative X-ray transmission through the sample to be measured. The sample was measured with a beam that had full-width half maxima of 26*×* 24 µm^2^ as measured by a knife-edge scan. The raster scan used a step size of 25 µm in both the vertical and horizontal directions, with continuous fly-scanning in the vertical direction. For each sample, a total of 220 projections over 6 different tilt angles were measured.

### 5.5 SAXS Tensor Tomography Reconstructions

The absorption data was used to align the projections with sub-pixel precision using cross-correlation algorithms [75]. The absorption tomogram was reconstructed using the Simultaneous Iterative Reconstruction Technique algortihm and subsequently segmented using Otsu’s method [73] to define the bone volume. The 2D scattering patterns obtained at each point were radially integrated to obtain 1D scattering curves in 32 azimuthal detector segments using PyFAI [76]. The data was radially binned in 2 different *q*-ranges, corresponding to the meridional collagen peak 0.08 - 0.115 nm^*−*1^ and the mineral scattering 0.13 - 1.9 nm^*−*1^. In the case of the collagen peak reconstruction, the scattering from the mineral background was subtracted in each azimuthal curve. Tensor tomographic reconstructions of the resulting data were performed with the python package *Mumott* [63] (Version 2.0). The scattering in each voxel was represented using a Gaussian kernel basis set. The optimization problem was defined by a squared loss function with Huber norm and total variation regularization, and was minimized by gradient descent. For each voxel, the mean intensity was calculated and the 3D RSM was approximated by a second order tensor, from which the main orientation and fractional anisotropy (FA) were calculated by eigenvector analysis. The FA is defined as:

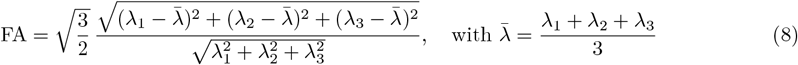

In their respective *q*-ranges, the largest eigenvector determined the main meridional collagen scattering orientation, whilst the smallest eigenvector determined the orientation perpendicular to the equatorial mineral particle scattering. The same reconstruction procedure was employed for *q*-resolved reconstructions (i.e. without radial binning) in both of the aformentioned scattering ranges, leading to full reconstruction of the reciprocal space within those ranges. These reconstructions were used to determine the meridional collagen D-spacing (*D* = 2*π/q*) and perform the mineral particle analysis on the spherically averaged intensity using the same methodology as described in section 5.3.

### 5.6 Statistical Analysis

Statistical analysis of the scanning SAXS/WAXS data was performed in Python using the *statsmodels* library [77]. Pairwise t-tests were conducted between sample medians from the right and left legs of the same donors, with anatomical location kept separate. For analysis of the full dataset, linear mixed-effects models were used. For each dependent variable *Y*, measured at pixel *i* from ROI *j* within sample *k* and donor *l*, the model was defined as:

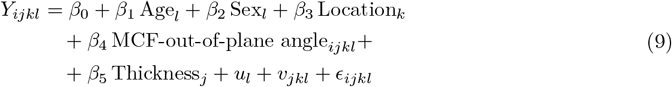

Here, *β*_0_ is the global intercept and *β*_1_–*β*_5_ are fixed-effect coefficients for age, sex, anatomical location, MCF out-of-plane angle and sample thickness, respectively. Thickness was included to account for variability arising from measurement uncertainty in sample thickness. The donor-specific random intercept 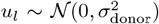 accounts for correlation among pixels from the same donor. The ROI random effect 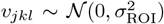 captures additional variance within each sample, with ROIs nested within samples. The residual term *ϵ*_*ijkl*_ *∼𝒩*(0, *σ*^2^) represents pixel-level variability within ROIs. Variance components were used to calculate intraclass correlation coefficients (ICCs) for donors and ROIs, defined as:

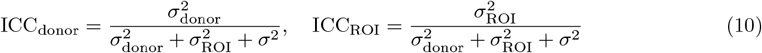

These ICCs quantify the proportion of total variance attributable to differences between donors and between ROIs within samples, respectively. All models were fitted using restricted maximum likelihood estimation (REML) in *statsmodels*. All p-values from the linear mixed-effects models were adjusted for multiple comparisons using the Benjamini-Hochberg procedure to control the false discovery rate.

## 6 Acknowledgments

This work was funded by the Swiss National Science Foundation (SNSF), grant number 200365. ML and AB acknowledge funding from the European research council (ERC-2020-StG 949301). MC has received funding from the European Unions Horizon 2020 research and innovation program under the Marie Skodowska-Curie grant agreement No 884104. Views and opinions expressed are those of the authors only and do not necessarily reflect those of the European Union or the European Research Council Executive Agency. Neither the European Union nor the granting authority can be held responsible for them. We acknowledge the Paul Scherrer Institut, Villigen, Switzerland for provision of synchrotron radiation beamtime at the beamline cSAXS of the SLS under proposal 20220327, and the MAX IV laboratory, Lund, Sweden, for provision of synchrotron radiation beamtime at the ForMAX beamline under proposal 20240337. The authors acknowledge the Statistical Consulting Group at ETH Zurich for providing advice during the statistical image analysis.

## 7 Author contributions

T.T.: Conceptualization, sample preparation, experiment, analysis, writing, figure creation. T.K.: Conceptualization, analysis support, reviewing and editing. A.B.: Experiment, analysis support, reviewing and editing. M.S.: Sample preparation, reviewing and editing. M.C.: Experiment, analysis support, reviewing and editing. S.F.B: local contact during synchrotron experiment at ForMAX (MAX IV), reviewing and editing. M.G-S: local contact during synchrotron experiment at cSAXS (SLS), reviewing and editing. P.Z: Conceptualization, supervision, funding acquisition, reviewing and editing. M.L.: Conceptualization, supervision, funding acquisition, experiment, analysis support, writing, reviewing and editing.

## References

[1] John A Kanis, Nicholas Norton, Nicholas C Harvey, Trolle Jacobson, Helena Johansson, Mattias Lorentzon, Eugene V McCloskey, Carl Willers, and Fredrik Borgström. SCOPE 2021: a new score-card for osteoporosis in Europe. Archives of Osteoporosis, 16(1):82, 2021.

[2] Carl Willers, Nicholas Norton, Nicholas C Harvey, Trolle Jacobson, Helena Johansson, Mattias Lorentzon, Eugene V McCloskey, Fredrik Borgström, and John A Kanis. Osteoporosis in Europe: a compendium of country-specific reports. Archives of Osteoporosis, 17(1):23, 2022.

[3] D. B. Burr. The contribution of the organic matrix to bone’s material properties. Bone, 31(1):8–11, 7 2002.

[4] Peter Fratzl and Richard Weinkamer. Nature’s hierarchical materials. Progress in Materials Science, 52(8):1263–1334, 11 2007.

[5] Elizabeth A. Zimmermann and Robert O. Ritchie. Bone as a Structural Material. Advanced Healthcare Materials, 4(9):1287–1304, 6 2015.

[6] Natalie Reznikov, Ron Shahar, and Steve Weiner. Bone hierarchical structure in three dimensions. Acta Biomaterialia, 10(9):3815–3826, 9 2014.

[7] P Zioupos. Ageing Human Bone: Factors Affecting its Biomechanical Properties and the Role of Collagen. Journal of Biomaterials Applications, 15(3):187–229, 1 2001.

[8] P. Roschger, P. Fratzl, J. Eschberger, and K. Klaushofer. Validation of quantitative backscattered electron imaging for the measurement of mineral density distribution in human bone biopsies. Bone, 23(4):319–326, 10 1998.

[9] R. B. Martin and J. Ishida. The relative effects of collagen fiber orientation, porosity, density, and mineralization on bone strength. Journal of biomechanics, 22(5):419–426, 1989.

[10] Tatiana Kochetkova, Cinzia Peruzzi, Oliver Braun, Jan Overbeck, Anjani K. Maurya, Antonia Neels, Michel Calame, Johann Michler, Philippe Zysset, and Jakob Schwiedrzik. Combining polarized Raman spectroscopy and micropillar compression to study microscale structure-property relationships in mineralized tissues. Acta Biomaterialia, 119:390–404, 1 2021.

[11] Ozan Akkus, Fran Adar, and Mitchell B. Schaffler. Age-related changes in physicochemical properties of mineral crystals are related to impaired mechanical function of cortical bone. Bone, 34(3):443–453, 3 2004.

[12] X. Wang, X. Shen, X. Li, and C. Mauli Agrawal. Age-related changes in the collagen network and toughness of bone. Bone, 31(1):1–7, 2002.

[13] E. P. Paschalis, D. N. Tatakis, S. Robins, P. Fratzl, I. Manjubala, R. Zoehrer, S. Gamsjaeger, B. Buchinger, A. Roschger, R. Phipps, A. L. Boskey, E. Dall’Ara, P. Varga, P. Zysset, K. Klaushofer, and P. Roschger. Lathyrism-induced alterations in collagen cross-links influence the mechanical properties of bone material without affecting the mineral. Bone, 49(6):1232–1241, 12 2011.

[14] D. Vashishth, G. J. Gibson, J. I. Khoury, M. B. Schaffler, J. Kimura, and D. P. Fyhrie. Influence of nonenzymatic glycation on biomechanical properties of cortical bone. Bone, 28(2):195–201, 2001.

[15] Ondřej Nikel, Atharva A. Poundarik, Stacyann Bailey, and Deepak Vashishth. Structural role of osteocalcin and osteopontin in energy dissipation in bone. Journal of biomechanics, 80:45–52, 10 2018.

[16] Mathilde Granke, Mark D. Does, and Jeffry S. Nyman. The Role of Water Compartments in the Material Properties of Cortical Bone. Calcified Tissue International, 97(3):292–307, 9 2015.

[17] M. B. Schaffler, K. Choi, and C. Milgrom. Aging and matrix microdamage accumulation in human compact bone. Bone, 17(6):521–525, 12 1995.

[18] P. Zioupos. Accumulation of in-vivo fatigue microdamage and its relation to biomechanical properties in ageing human cortical bone. Journal of Microscopy, 201(2):270–278, 2001.

[19] Jakob Schwiedrzik, Rejin Raghavan, Alexander Bürki, Victor LeNader, Uwe Wolfram, Johann Michler, and Philippe Zysset. In situ micropillar compression reveals superior strength and ductility but an absence of damage in lamellar bone. Nature Materials, 13(7):740–747, 2014.

[20] Ottman A. Tertuliano and Julia R. Greer. The nanocomposite nature of bone drives its strength and damage resistance. Nature materials, 15(11):1195–1202, 11 2016.

[21] Alexander Groetsch, Aurélien Gourrier, Jakob Schwiedrzik, Michael Sztucki, Rainer J. Beck, Jonathan D. Shephard, Johann Michler, Philippe K. Zysset, and Uwe Wolfram. Compressive behaviour of uniaxially aligned individual mineralised collagen fibres at the micro- and nanoscale. Acta Biomaterialia, 89:313–329, 4 2019.

[22] Daniele Casari, Johann Michler, Philippe Zysset, and Jakob Schwiedrzik. Microtensile properties and failure mechanisms of cortical bone at the lamellar level. Acta Biomaterialia, 120:135–145, 1 2020.

[23] Fei Hang and Asa H. Barber. Nano-mechanical properties of individual mineralized collagen fibrils from bone tissue. Journal of the Royal Society Interface, 8(57):500, 4 2010.

[24] Ines Jimenez-Palomar, Anna Shipov, Ron Shahar, and Asa H. Barber. Structural orientation dependent sub-lamellar bone mechanics. Journal of the Mechanical Behavior of Biomedical Materials, 52:63–71, 12 2015.

[25] K. E. Rudman, R. M. Aspden, and J. R. Meakin. Compression or tension? The stress distribution in the proximal femur. BioMedical Engineering Online, 5(1):12–, 2 2006.

[26] Peter M De Bakker, Sarah L Manske, Vincent Ebacher, Thomas R Oxland, Peter A Cripton, and Pierre Guy. During sideways falls proximal femur fractures initiate in the superolateral cortex: Evidence from high-speed video of simulated fractures. Journal of Biomechanics, 42(12):1917–1925, 8 2009.

[27] Shashank Nawathe, Hosna Akhlaghpour, Mary L. Bouxsein, and Tony M. Keaveny. Microstructural failure mechanisms in the human proximal femur for sideways fall loading. Journal of bone and mineral research, 29(2):507–515, 2 2014.

[28] Paul M. Mayhew, C. David Thomas, John G. Clement, Nigel Loveridge, Thomas J. Beck, William Bonfield, Chris J. Burgoyne, and Jonathan Reeve. Relation between age, femoral neck cortical stability, and hip fracture risk. Lancet, 366(9480):129–135, 7 2005.

[29] K. L. Bell, N. Loveridge, J. Power, N. Garrahan, B. F. Meggitt, and J. Reeve. Regional differences in cortical porosity in the fractured femoral neck. Bone, 24(1):57–64, 1 1999.

[30] H. Blain, Pascale Chavassieux, N. Portero-Muzy, F. Bonnel, F. Canovas, M. Chammas, P. Maury, and P. D. Delmas. Cortical and trabecular bone distribution in the femoral neck in osteoporosis and osteoarthritis. Bone, 43(5):862–868, 11 2008.

[31] Marios Georgiadis, Ralph Müller, and Philipp Schneider. Techniques to assess bone ultrastructure organization: Orientation and arrangement of mineralized collagen fibrils. Journal of the Royal Society Interface, 13(119), 6 2016.

[32] Dakota M. Binkley and Kathryn Grandfield. Advances in Multiscale Characterization Techniques of Bone and Biomaterials Interfaces. ACS biomaterials science & engineering, 4(11):3678–3690, 11 2018.

[33] P Fratzl, S Schreiber, and K Klaushofer. Bone mineralization as studied by small-angle x-ray scattering. Connective Tissue Research, 34(4):247–254, 1996.

[34] Peter Fratzl, Himadri S. Gupta, Oskar Paris, Angelika Valenta, Paul Roschger, and Klaus Klaushofer. Diffracting “stacks of cards” - some thoughts about small-angle scattering from bone. Progress in Colloid and Polymer Science, 130:33–39, 2005.

[35] Silvia Pabisch, Wolfgang Wagermaier, Thomas Zander, Chenghao Li, and Peter Fratzl. Imaging the Nanostructure of Bone and Dentin Through Small- and Wide-Angle X-Ray Scattering. Methods in Enzymology, 532:391–413, 1 2013.

[36] Mathias H. Bünger, Hans Oxlund, Toke K. Hansen, Søren Sørensen, Bo M. Bibby, Jesper S. Thomsen, Bente L. Langdahl, Flemming Besenbacher, Jan S. Pedersen, and Henrik Birkedal. Strontium and bone nanostructure in normal and ovariectomized rats investigated by scanning small-angle X-ray scattering. Calcified tissue international, 86(4):294–306, 4 2010.

[37] C Giannini, D Siliqi, M Ladisa, D Altamura, A Diaz, A Beraudi, T Sibillano, L De Caro, S Stea, F Baruffaldi, and O Bunk. Scanning SAXS-WAXS microscopy on osteoarthritis-affected bone-an age-related study. J. Appl. Cryst, 47:110–117, 2014.

[38] Heikki Suhonen, Manuel Fernández, Ritva Serimaa, and Pekka Suortti. Simulation of small-angle x-ray scattering from collagen fibrils and comparison with experimental patterns. Physics in Medicine & Biology, 50(22):5401, 11 2005.

[39] Richard S. Bear. X-Ray Diffraction Studies on Protein Fibers. I. The Large Fiber-Axis Period of Collagen. Journal of the American Chemical Society, 66(8):1297–1305, 8 1944.

[40] Stephen W. White, David J.S. Hulmes, Andrew Miller, and Peter A. Timmins. Collagen–mineral axial relationship in calcified turkey leg tendon by X-ray and neutron diffraction. Nature, 266(5601):421–425, 3 1977.

[41] Aurélien Gourrier, Chenghao Li, Stefan Siegel, Oskar Paris, Paul Roschger, Klaus Klaushofer, and Peter Fratzl. Scanning small-angle X-ray scattering analysis of the size and organization of the mineral nanoparticles in fluorotic bone using a stack of cards model. J. Appl. Cryst, 43:1385–1392, 2010.

[42] P. Fratzl, H. S. Gupta, E. P. Paschalis, and P. Roschger. Structure and mechanical quality of the collagen–mineral nano-composite in bone. Journal of Materials Chemistry, 14(14):2115–2123, 7 2004.

[43] O Bunk, M Bech, H H Jensen, R Feidenhansy, T Binderup, A Menzel, and F Pfeiffer. Multimodal x-ray scatter imaging. New Journal of Physics, 11(12):123016, 12 2009.

[44] N. Fratzl-Zelman, P. Roschger, A. Gourrier, M. Weber, B. M. Misof, N. Loveridge, J. Reeve, K. Klaushofer, and P. Fratzl. Combination of nanoindentation and quantitative backscattered electron imaging revealed altered bone material properties associated with femoral neck fragility. Calcified Tissue International, 85(4):335–343, 10 2009.

[45] Tengteng Tang, Wolfgang Wagermaier, Roman Schuetz, Qiong Wang, Felipe Eltit, Peter Fratzl, and Rizhi Wang. Hypermineralization in the femoral neck of the elderly. Acta Biomaterialia, 89:330–342, 4 2019.

[46] Marianne Liebi, Marios Georgiadis, Andreas Menzel, Philipp Schneider, Joachim Kohlbrecher, Oliver Bunk, and Manuel Guizar-Sicairos. Nanostructure surveys of macroscopic specimens by small-angle scattering tensor tomography. Nature, 527(7578):349–352, 11 2015.

[47] Tilman A. Grünewald, Marianne Liebi, Nina K. Wittig, Andreas Johannes, Tanja Sikjaer, Lars Rejnmark, Zirui Gao, Martin Rosenthal, Manuel Guizar-Sicairos, Henrik Birkedal, and Manfred Burghammer. Mapping the 3D orientation of nanocrystals and nanostructures in human bone: Indications of novel structural features. Science Advances, 6(24):4171–4183, 6 2020.

[48] Tilman A. Grünewald, Andreas Johannes, Nina K. Wittig, Jonas Palle, Alexander Rack, Manfred Burghammer, and Henrik Birkedal. Bone mineral properties and 3D orientation of human lamellar bone around cement lines and the Haversian system. IUCrJ, 10(Pt 2):189–198, 2 2023.

[49] Leonard C. Nielsen, Paul Erhart, Manuel Guizar-Sicairos, and Marianne Liebi. Small-angle scattering tensor tomography algorithm for robust reconstruction of complex textures. Acta Crystallographica Section A: Foundations and Advances, 79(Pt 6):515–526, 10 2023.

[50] Marios Georgiadis, Manuel Guizar-Sicairos, Oliver Gschwend, Peter Hangartner, Oliver Bunk, Ralph Möller, and Philipp Schneider. Ultrastructure Organization of Human Trabeculae Assessed by 3D sSAXS and Relation to Bone Microarchitecture. PLOS ONE, 11(8), 8 2016.

[51] Paolino De Falco, Richard Weinkamer, Wolfgang Wagermaier, Chenghao Li, Tim Snow, Nicholas J Terrill, Himadri S Gupta, Pawan Goyal, Martin Stoll, Peter Benner, and Peter Fratzl. Tomographic X-ray scattering based on invariant reconstruction: analysis of the 3D nanostructure of bovine bone. J. Appl. Cryst, 54:486–497, 2021.

[52] Fabian Pedregosa, Gaël Varoquaux, Alexandre Gramfort, Vincent Michel, Bertrand Thirion, Olivier Grisel, Mathieu Blondel, Peter Prettenhofer, Ron Weiss, Vincent Dubourg, Jake Vanderplas, Alexandre Passos, Matthieu Brucher, Matthieu Perrot, and Édouard Duchesnay. Scikit-learn: Machine Learning in Python. Journal of Machine Learning Research, 12(85):2825–2830, 2011.

[53] Nigel Loveridge, Jon Power, Jonathan Reeve, and Alan Boyde. Bone mineralization density and femoral neck fragility. Bone, 35(4):929–941, 10 2004.

[54] Simon von Kroge, Julian Stürznickel, Ulrich Bechler, Kilian Elia Stockhausen, Julian Eissele, Jan Hubert, Michael Amling, Frank Timo Beil, Björn Busse, and Tim Rolvien. Impaired bone quality in the superolateral femoral neck occurs independent of hip geometry and bone mineral density. Acta Biomaterialia, 141:233–243, 3 2022.

[55] W. Wagermaier, H. S. Gupta, A. Gourrier, M. Burghammer, P. Roschger, and P. Fratzl. Spiral twisting of fiber orientation inside bone lamellae. Biointerphases, 1(1):1–5, 3 2006.

[56] M Granke, A Gourrier, F Rupin, K Raum, and F Peyrin. Microfibril Orientation Dominates the Microelastic Properties of Human Bone Tissue at the Lamellar Length Scale. PLOS ONE, 8(3), 2013.

[57] Maria Grazia Ascenzi, Antonio Ascenzi, Alessandro Benvenuti, Manfred Burghammer, Silvia Panzavolta, and Adriana Bigi. Structural differences between “dark” and “bright” isolated human osteonic lamellae. Journal of Structural Biology, 141(1):22–33, 1 2003.

[58] Joseph P.R.O. Orgel, Thomas C. Irving, Andrew Miller, and Tim J. Wess. Microfibrillar structure of type I collagen in situ. Proceedings of the National Academy of Sciences of the United States of America, 103(24):9001–9005, 6 2006.

[59] T. J. Wess. Collagen Fibrillar Structure and Hierarchies. In Collagen: Structure and Mechanics, pages 49–80. Springer, Boston, MA, 2008.

[60] P. Roschger, B. M. Grabner, S. Rinnerthaler, W. Tesch, M. Kneissel, A. Berzlanovich, K. Klaushofer, and P. Fratzl. Structural development of the mineralized tissue in the human L4 vertebral body. Journal of structural biology, 136(2):126–136, 2001.

[61] J. Power, N. Loveridge, N. Rushton, M. Parker, and J. Reeve. Evidence for bone formation on the external “periosteal” surface of the femoral neck: A comparison of intracapsular hip fracture cases and controls. Osteoporosis International, 14(2):141–145, 2003.

[62] Petar Milovanovic and Björn Busse. Inter-site Variability of the Human Osteocyte Lacunar Network: Implications for Bone Quality. Current Osteoporosis Reports, 17(3):105–115, 6 2019.

[63] Leonard C. Nielsen, Mads Carlsen, Sici Wang, Arthur Baroni, Torne Tänzer, Marianne Liebi, and Paul Erhart. MUMOTT: A Python package for the analysis of multi-modal tensor tomography data. Journal of Applied Crystallography, 58(Pt 5):1834–1845, 10 2025.

[64] Marianne Liebi, Marios Georgiadis, Joachim Kohlbrecher, Mirko Holler, Jörg Raabe, Ivan Usov, Andreas Menzel, Philipp Schneider, Oliver Bunk, and Manuel Guizar-Sicairos. Small-angle X-ray scattering tensor tomography: model of the three-dimensional reciprocal-space map, reconstruction algorithm and angular sampling requirements. Acta Crystallographica Section A: Foundations and Advances, 74(1):12–24, 1 2018.

[65] Marios Georgiadis, Manuel Guizar-Sicairos, Alexander Zwahlen, Andreas J. Trüssel, Oliver Bunk, Ralph Müller, and Philipp Schneider. 3D scanning SAXS: A novel method for the assessment of bone ultrastructure orientation. Bone, 71:42–52, 2 2015.

[66] I. Zizak, P. Roschger, O. Paris, B. M. Misof, A. Berzlanovich, S. Bernstorff, H. Amenitsch, K. Klaushofer, and P. Fratzl. Characteristics of mineral particles in the human bone/cartilage interface. Journal of Structural Biology, 141(3):208–217, 3 2003.

[67] M. H. Bünger, Morten Foss, K. Erlacher, H. Li, X. Zou, B. L. Langdahl, C. Bünger, H. Birkedal, F. Besenbacher, and J. S. Pedersen. Bone nanostructure near titanium and porous tantalum implants studied by scanning small angle x-ray scattering. European cells & materials, 12:81–91, 2006.

[68] Mathias Hauge Bünger, Morten Foss, Kurt Erlacher, Mads Bruun Hovgaard, Jacques Chevallier, Bente Langdahl, Cody Bünger, Henrik Birkedal, Flemming Besenbacher, and Jan Skov Pedersen. Nanostructure of the neurocentral growth plate: Insight from scanning small angle X-ray scattering, atomic force microscopy and scanning electron microscopy. Bone, 39(3):530–541, 2006.

[69] Shoaib Attar, Hanna Isaksson, Lars Jehpsson, Bjorn E. Rosengren, Magnus K. Karlsson, and Lorenzo Grassi. Variations over time in proximal femoral strength in young adult men are not explained by areal bone mineral density alone. JBMR plus, 9(10), 10 2025.

[70] Benjamin Voumard, Pia Stefanek, Michael Pretterklieber, Dieter Pahr, and Philippe Zysset. Influence of aging on mechanical properties of the femoral neck using an inverse method. Bone Reports, 17:101638, 12 2022.

[71] Mirko Holler, Johannes Ihli, Esther H.R. Tsai, Fabio Nudelman, Mariana Verezhak, Wilma D.J. Van De Berg, and Sarah H. Shahmoradian. A lathe system for micrometre-sized cylindrical sample preparation at room and cryogenic temperatures. Journal of Synchrotron Radiation, 27(2):472–476, 3 2020.

[72] Jérôme Kieffer, Julien Orlans, Nicolas Coquelle, Samuel Debionne, Shibom Basu, Alejandro Homs, Gianluca Santoni, and Daniele De Sanctis. Application of signal separation to diffraction image compression and serial crystallography. urn:issn:1600-5767, 58(1):138–153, 2 2025.

[73] Nobuyuki Otsu. A Threshold Selection Method from Gray-Level Histograms. IEEE Trans Syst Man Cybern, SMC-9(1):62–66, 1979.

[74] Andre Guinier and Gérard Fournet. Small-Angle Scattering of X-Rays. John Wiley and Sons, New York, 1955.

[75] Manuel Guizar-Sicairos, Samuel T Thurman, and James R Fienup. Efficient subpixel image registration algorithms. Optics Letters, 100(2), 2008.

[76] Giannis Ashiotis, Aurore Deschildre, Zubair Nawaz, Jonathan P. Wright, Dimitrios Karkoulis, Frédéric Emmanuel Picca, and Jérôme Kieffer. The fast azimuthal integration Python library: pyFAI. Journal of Applied Crystallography, 48(2):510–519, 3 2015.

[77] Skipper Seabold and Josef Perktold. Statsmodels: Econometric and Statistical Modeling with Python. In 9th Python in Science Conference, pages 57–61, Austin, 6 2010.

