## Supporting Information for "Combination of 3D and 2D small and wide angle X-ray scattering imaging reveals diminished bone quality in the superior human femoral neck cortex"

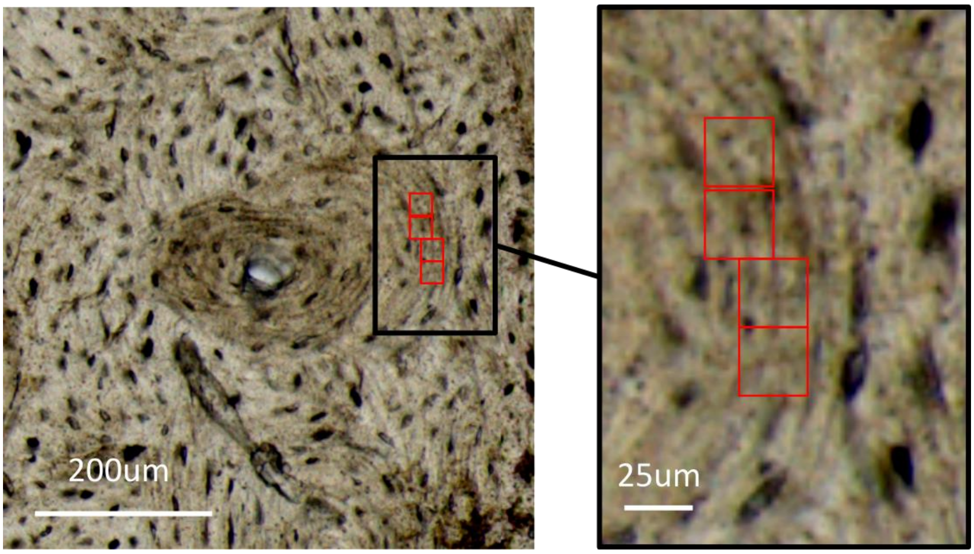


*Figure S1: Representation of beamsize/voxel size of 25µm² compared to the bone microstructure. The voxel corresponds to a size in which lamellae are predominantly aligned in parallel, exhibiting minimal curvature over this length scale, leading to non-axially symmetric fibre distributions in the transversal plane.*

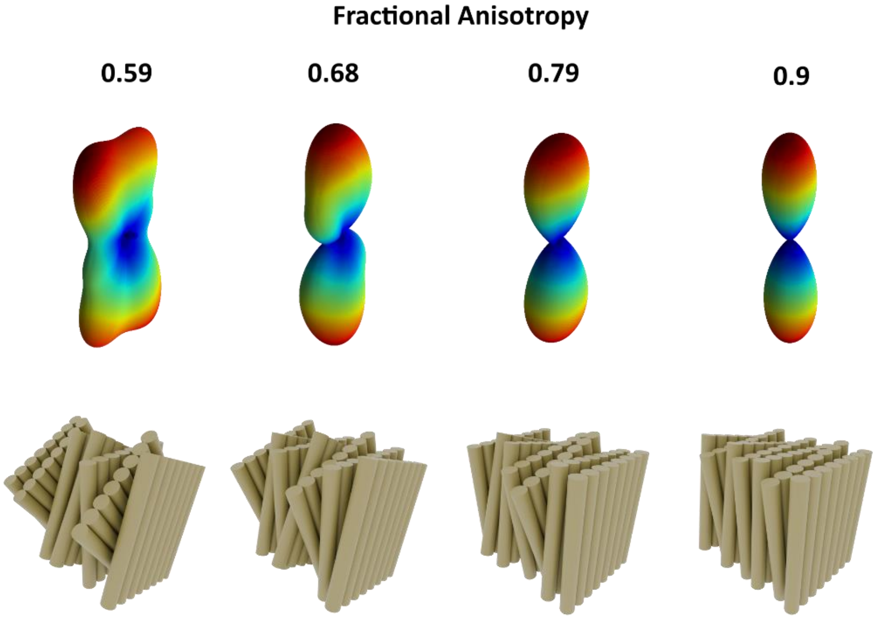


*Figure S2: Representative meridional collagen RSM's of different 3D Fractional Anisotropies extracted from the inferior side of sample pair 2 in relation to the possible underlying lamellar structure*

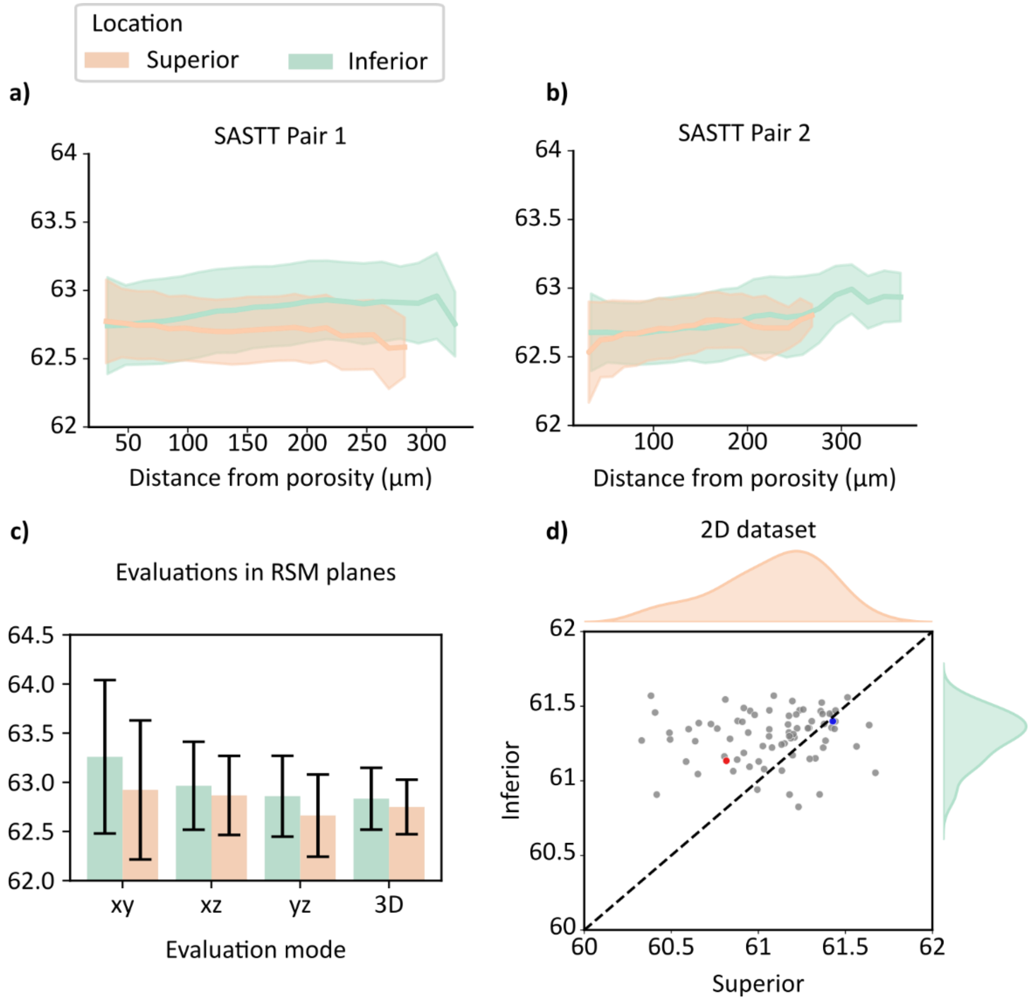


*Figure S3: (a,b) Collagen D-spacing (nm) in the SASTT sample pairs, in function of distance from porosity. In pair 1, the collagen D-spacing is slightly lower on the superior side, while they are similar in pair 2. (c) Evaluation of collagen D-spacing in 3 orthogonal planes oriented w.r.t the bone axis z. No significant differences were noted.(d) Scatter plot of the mean D-spacing of all femoral necks (x-axis: superior, y-axis: inferior). There is a significant but very small (0.3%) decrease in the average on the superior side. The blue and red points indicate the sample pairs from figure 4.*

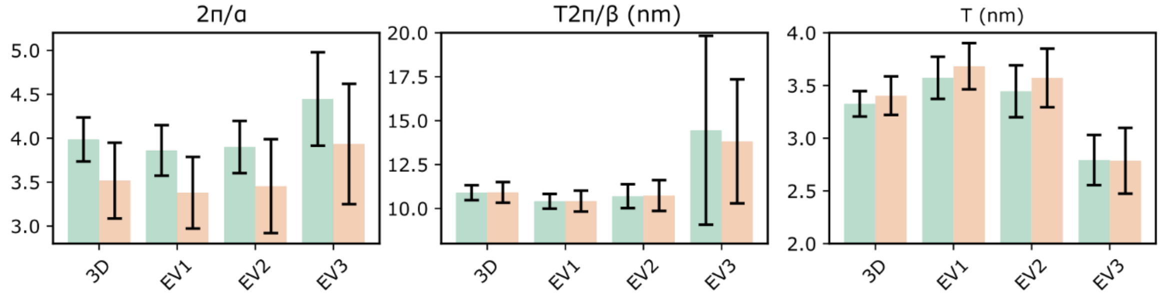


*Figure S4: Mineral size and arrangement parameters calculated along the Eigenvectors of the Mineral scattering signal ($q = 0.18 – 0.5 nm^{-1}$). The smallest eigenvector (EV3) indicates the direction of the weakest mineral scattering, displaying short-range order ($2\pi/\alpha$) and inter-platelet distance ($\frac{T2\pi}{\beta}$) values up to 18% and 38% higher than in the strongest scattering direction (EV1). The T-parameter is up to 25% lower in the weakest scattering directions than in the strongest. These values display the extent of the 3D directional differences in the mineral size and arrangement at the 25μmlength scale.*

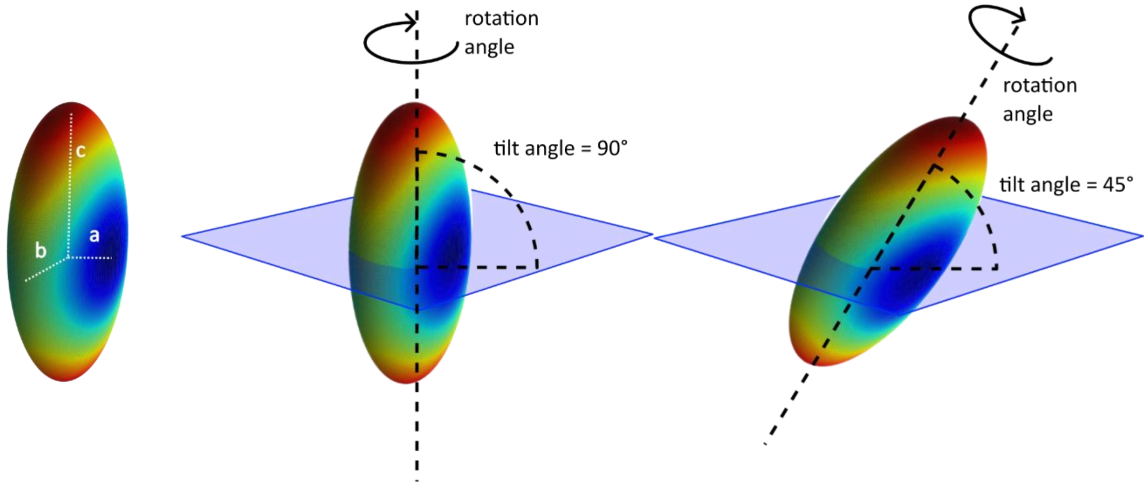


*Figure S5: Representation of a generalized ellipsoid with half axes a,b,c, with the out-of-plane angle and transversal measurement plane (blue plane).*

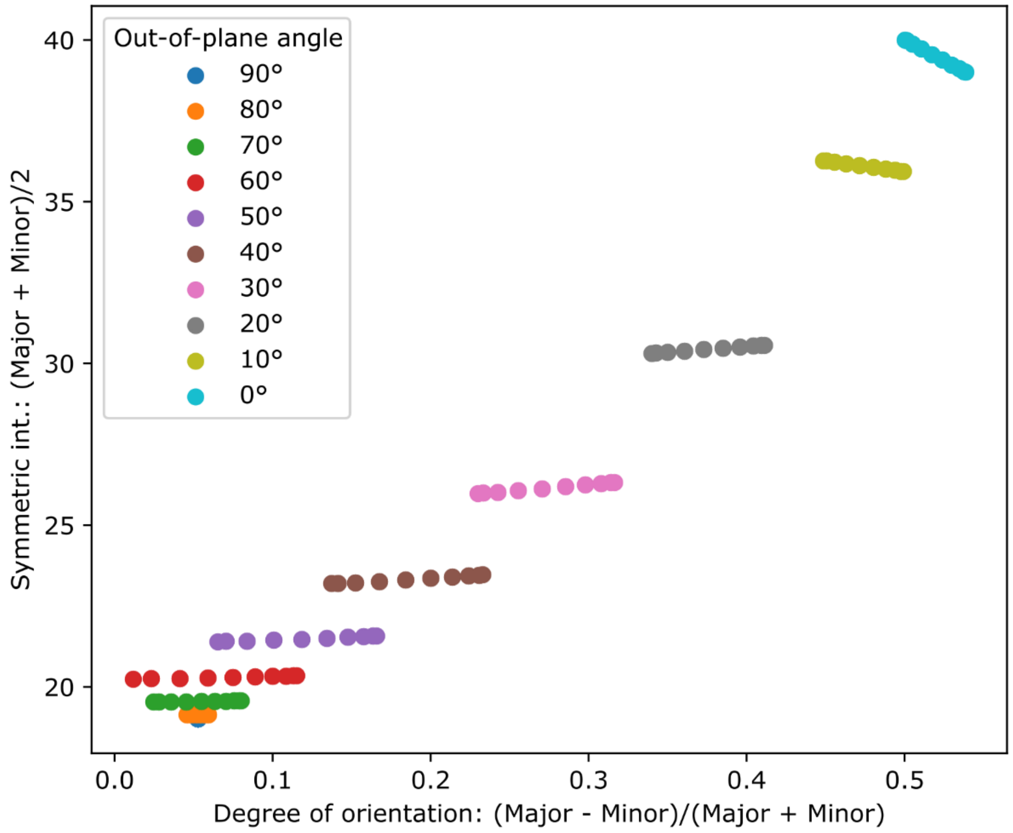


*Figure S6: Scatter plot of mean intensity vs measured degree of orientation in the transversal plane for a generalized ellipsoid (a = 18, b=20, c=60) tilted and rotated at various angles. Each color corresponds to a tilt (out-of-plane) angle, and the spread within a tilt angle is due to the transversal anisotropy (a and b axes). The out-of-plane angle can theoretically be determined from the combined measurement of the mean intensity and the degree of orientation in the transversal plane for this case.*

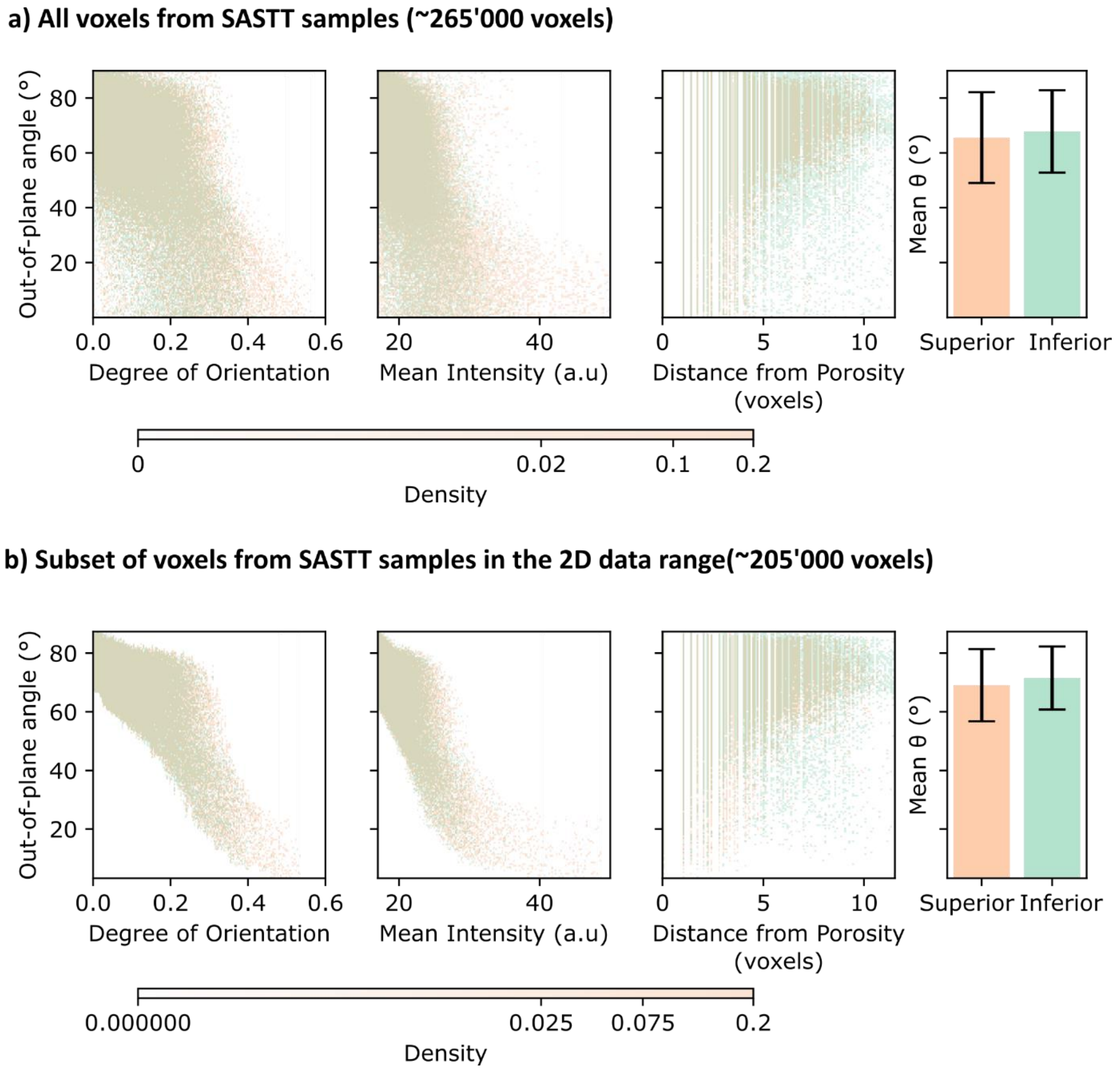


*Figure S7: Correlations between out-of-plane angle of meridional collagen scattering with the model parameters, by anatomical location: Degree of orientation and mean intensity of the transversal slices, distance of the voxel from porosity and anatomical location of the voxel. (a) All voxels from the 4 SASTT samples. (b) Subset of voxels in the 2D data range (between the 5$^{th}$ and 95$^{th}$ percentiles of mean intensity for a given 2D degree of orientation, see Figure 3d). The individual correlations of the out-of-plane angle with the degree of orientation and mean intensities are broad, but can be combined to obtain a stronger correlation with the out-of-plane angle (Figure 3c)*

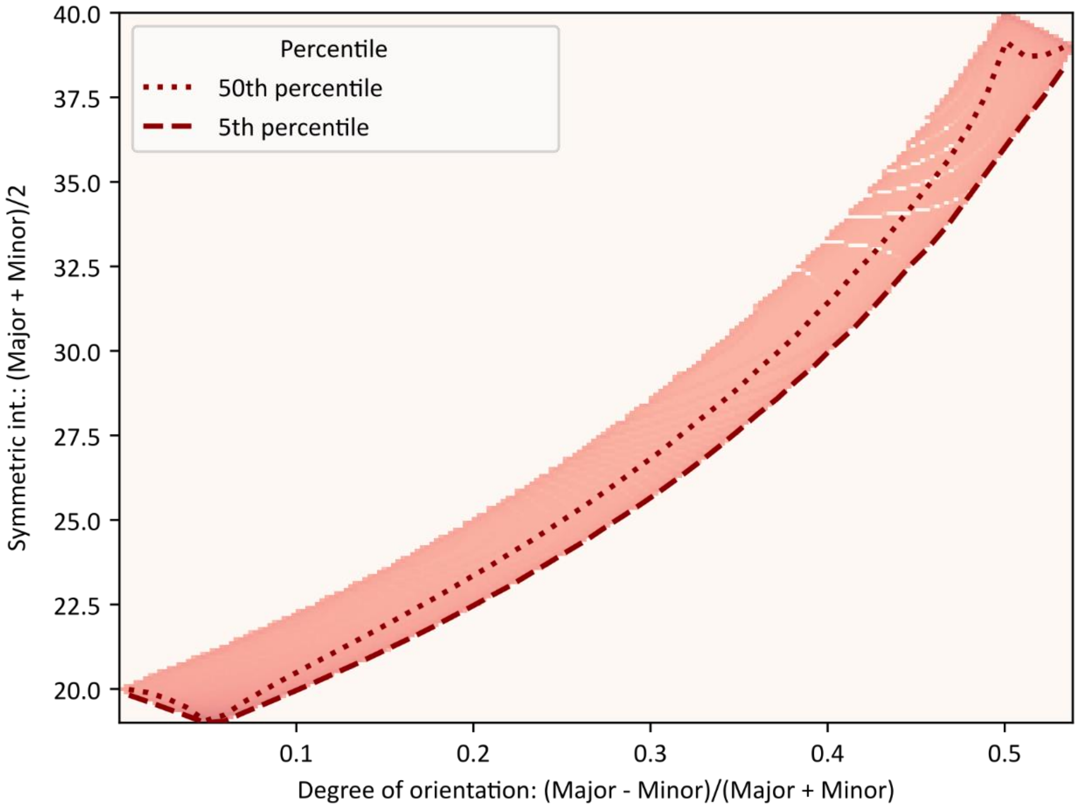


*Figure S8: Density plot of mean intensity vs measured degree of orientation in the transversal plane for a generalized ellipsoid (a = 18, b=20, c=60) tilted and rotated at various angles. The 50$^{th}$ percentile line in each degree of orientation bin illustrates the asymmetric mean intensity distribution obtained, which correlates to the distribution found in the experimental*

*data.*

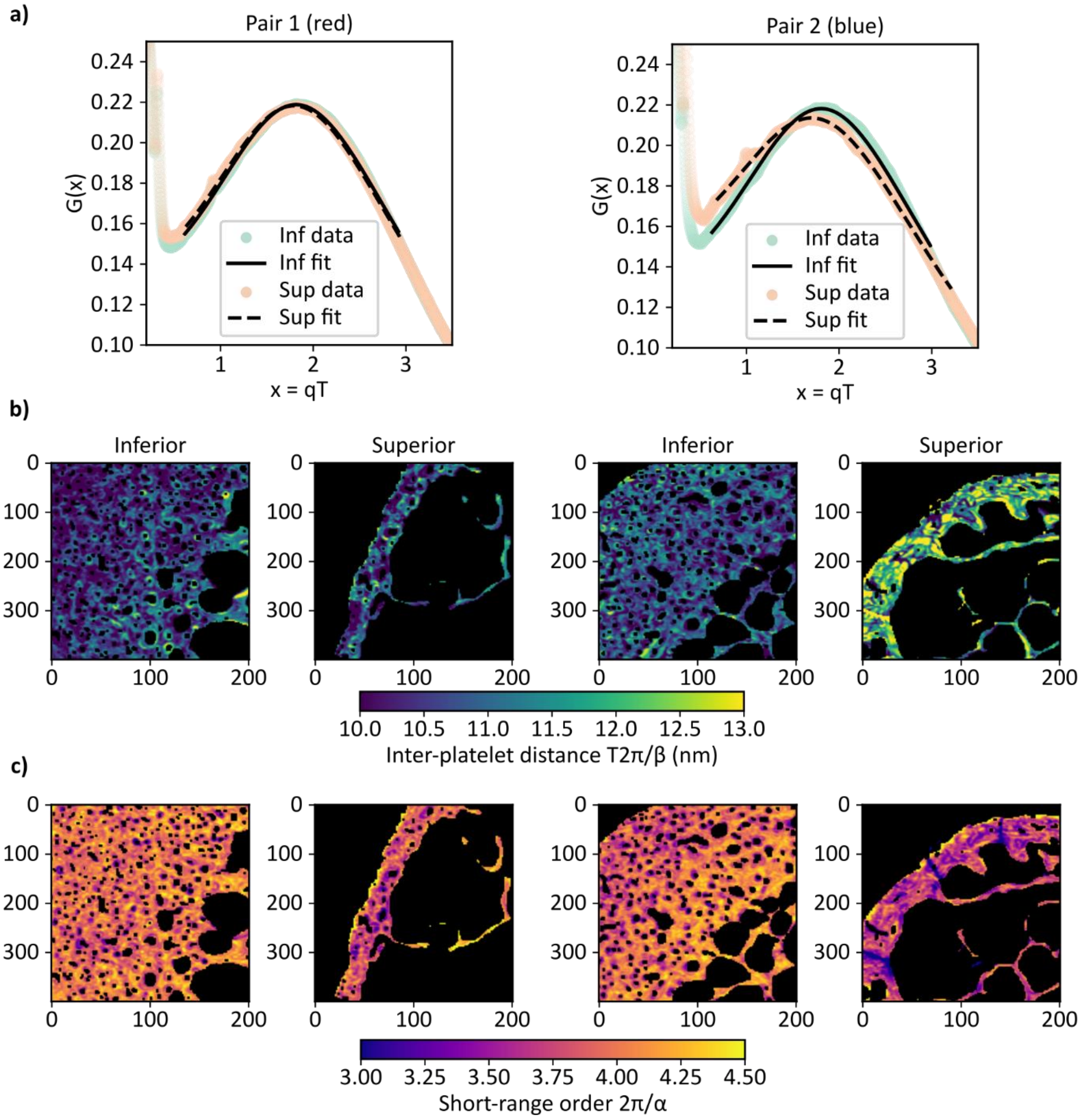


*Figure S9: Mineral arrangement parameters for the example pairs shown in figure 4. a) Mean G(x) curves and the corresponding fits using the stack-of-cards model. b) interplatelet-distance parameter c) short range order parameter.*

*Table S1: Variance decomposition of absorption and orientation parameters in the linear mixed-effects model*

| | X-ray absorption | Mer. Collagen Mean Intensity | Mer. Collagen Degree of Orientation | Equ. Collagen Mean Intensity | Equ. Collagen Degree of Orientaion | Mineral Mean Intensity | Mineral Degree of Orientation | HA (002) Mean Intensity | HA (002) Degree of Orientation | MCF out-of-plane angle |
|---|---|---|---|---|---|---|---|---|---|---|
| **ICC_Donor (%)** | 4.27 | 4.49 | 2.74 | 8.68 | 3.27 | 10.65 | 4.57 | 14.26 | 1.77 | 2.36 |
| **ICC_ROI (%)** | 18.30 | 11.63 | 10.31 | 16.82 | 10.97 | 13.29 | 8.48 | 11.66 | 8.40 | 17.35 |
| **Residual_ Fraction (%)** | 77.42 | 83.88 | 86.95 | 74.50 | 85.75 | 76.06 | 86.95 | 74.08 | 89.83 | 80.28 |

*Table S2: Varaince decomposition of structural parameters in the linear mixed-effects models*

| | T-paramenter | Mineral disorder | Inter-platelet distance | Mer. Collagen D-spacing | Equ. Collagen D-spacing | Collagen ratio (Mer. Vs Equ.) | Crystal Size | HA (002) D-spacing | Alpha | Beta |
|---|---|---|---|---|---|---|---|---|---|---|
| **ICC_Donor (%)** | 7.31 | 20.06 | 8.18 | 1.07 | 3.46 | 6.40 | 4.50 | 17.11 | 19.90 | 12.85 |
| **ICC_ROI (%)** | 13.83 | 9.74 | 12.45 | 4.35 | 18.71 | 13.64 | 4.74 | 6.51 | 9.94 | 10.61 |
| **Residual_ Fraction (%)** | 78.86 | 70.20 | 79.37 | 94.59 | 77.83 | 79.95 | 90.76 | 76.38 | 70.16 | 76.54 |

*Table S3: Results from the linear mixed-effects model for all the analyzed parameters. The coefficients and p-values are color-coded by their level of significance ($p < 0.001$, $p<0.01$, $p<0.05$, $p>0.05$). All p-values are adjusted for multiple comparisons using the Benjamini-Hochberg procedure to control the false discovery rate.*

| | | **Age** | | **Sex[M]** | | **Location [Sup]** | | **MCF Out-of-plane angle** | | **Thickness** | |
|---|---|---|---|---|---|---|---|---|---|---|---|
| **Parameter** | **Intercept $\beta_0$** | **Coeficient $\beta_1$** | **p-value** | **Coeficient $\beta_2$** | **p-value** | **Coeficient $\beta_3$** | **p-value** | **Coeficient $\beta_4$ (per 10°)** | **p-value** | **Coeficient $\beta_5$ (per $\mu m$)** | **p-value** |
| X-ray absorption ($cm^{-1}$) | 9.48e+00 | 7.28e-03 | 0.914 | -4.40e-02 | 0.953 | -1.02e+00 | <1e-16 | 2.63e-01 | <1e-16 | 2.96e-02 | <1e-16 |
| Coll. Mer. Mean int. | 2.82e+01 | -7.92e-04 | 0.914 | 1.41e-02 | 0.953 | 1.94e-01 | <1e-16 | -1.44e+00 | <1e-16 | 4.93e-04 | <0.001 |
| Coll. Mer. DOO | 3.85e-01 | -1.06e-05 | 0.914 | 2.84e-04 | 0.953 | 6.64e-03 | <1e-16 | -4.80e-02 | <1e-16 | 1.29e-04 | <1e-16 |
| Coll. Equ. Mean int. | 1.61e+01 | -4.19e-03 | 0.914 | 6.37e-02 | 0.736 | -5.91e-01 | <1e-16 | 3.97e-01 | <1e-16 | 1.28e-02 | <1e-16 |
| Coll. Equ. DOO | 4.34e-02 | 2.86e-05 | 0.914 | -8.11e-04 | 0.736 | 6.02e-03 | <1e-16 | -3.22e-03 | <1e-16 | 9.56e-07 | 0.604 |
| Mineral Mean int. | -6.68e+02 | -1.92e+00 | 0.914 | 1.77e+00 | 0.956 | -7.83e+01 | <1e-16 | 2.09e+02 | <1e-16 | 3.89e+00 | <1e-16 |
| Mineral DOO | 6.10e-01 | -2.26e-04 | 0.914 | 3.80e-03 | 0.872 | 4.91e-02 | <1e-16 | -6.34e-02 | <1e-16 | 1.67e-04 | <1e-16 |
| HA (002) Mean int. | 3.29e+02 | 4.93e-02 | 0.914 | -4.55e+00 | 0.736 | -5.21e-01 | <0.001 | -2.43e+01 | <1e-16 | -3.15e-01 | <1e-16 |
| HA (002) DOO | 6.55e-01 | 2.86e-04 | 0.914 | 6.80e-03 | 0.736 | 1.99e-02 | <1e-16 | -7.17e-02 | <1e-16 | 1.11e-04 | 0.001 |
| MCF out-of-plane angle | 72.43e+00 | 2.10e-02 | 0.914 | -6.97e-01 | 0.736 | -2.35e+00 | <1e-16 | N/A | N/A | 5.00e-02 | <1e-16 |
| Power law exp. | 1.56e+00 | -4.25e-04 | 0.914 | -1.41e-02 | 0.378 | 8.00e-02 | <1e-16 | 1.14e-02 | <1e-16 | -3.83e-05 | 0.007 |
| T-parameter (nm) | 2.22e+00 | 1.54e-03 | 0.914 | 5.07e-02 | 0.953 | 1.44e-01 | <1e-16 | 7.83e-02 | <1e-16 | -2.43e-04 | <0.001 |
| Crystal Size (nm) | 7.54e+01 | -1.49e-02 | 0.914 | -2.48e-01 | 0.953 | 1.25e+00 | <1e-16 | -3.80e+00 | <1e-16 | 1.34e-01 | <1e-16 |
| Coll. Mer. D-spacing (nm) | 5.81e+01 | 6.85e-04 | 0.914 | -1.80e-02 | 0.736 | -4.25e-01 | <1e-16 | 4.41e-01 | <1e-16 | 6.36e-03 | <1e-16 |
| HA (002) D-spacing (nm) | 3.43e-01 | -1.32e-05 | 0.914 | 1.31e-04 | 0.736 | 2.49e-04 | <1e-16 | 2.97e-04 | <1e-16 | 1.21e-05 | <1e-16 |
| Coll. Equ. D-spacing (nm) | 7.57e+01 | -6.82e-03 | 0.914 | -4.52e-01 | 0.953 | -1.24e+00 | <1e-16 | 1.44e+00 | <1e-16 | 1.57e-02 | <1e-16 |
| Coll. ratio (Mer. vs Equ.) | 1.65e+00 | 9.99e-05 | 0.914 | -2.42e-03 | 0.736 | 4.06e-02 | <1e-16 | -9.72e-02 | <1e-16 | -6.72e-04 | <1e-16 |
| Alpha | 1.56e+00 | -4.12e-04 | 0.914 | -1.66e-02 | 0.736 | 7.91e-02 | <1e-16 | 9.14e-03 | <1e-16 | -2.02e-04 | <1e-16 |
| Beta | 1.59e+00 | -2.04e-04 | 0.914 | -7.81e-03 | 0.736 | 1.91e-02 | <1e-16 | 1.58e-02 | <1e-16 | -1.17e-04 | <1e-16 |
| Mineral Disorder | 4.05e+00 | 8.00e-04 | 0.914 | 4.03e-02 | 0.087 | -1.86e-01 | <1e-16 | -2.06e-02 | <1e-16 | 4.90e-04 | <1e-16 |
| Inter-platelet Distance (nm) | 8.97e+00 | 6.96e-03 | 0.914 | 2.45e-01 | 0.953 | 4.48e-01 | <1e-16 | -1.79e-01 | <1e-16 | 3.53e-04 | 0.001 |

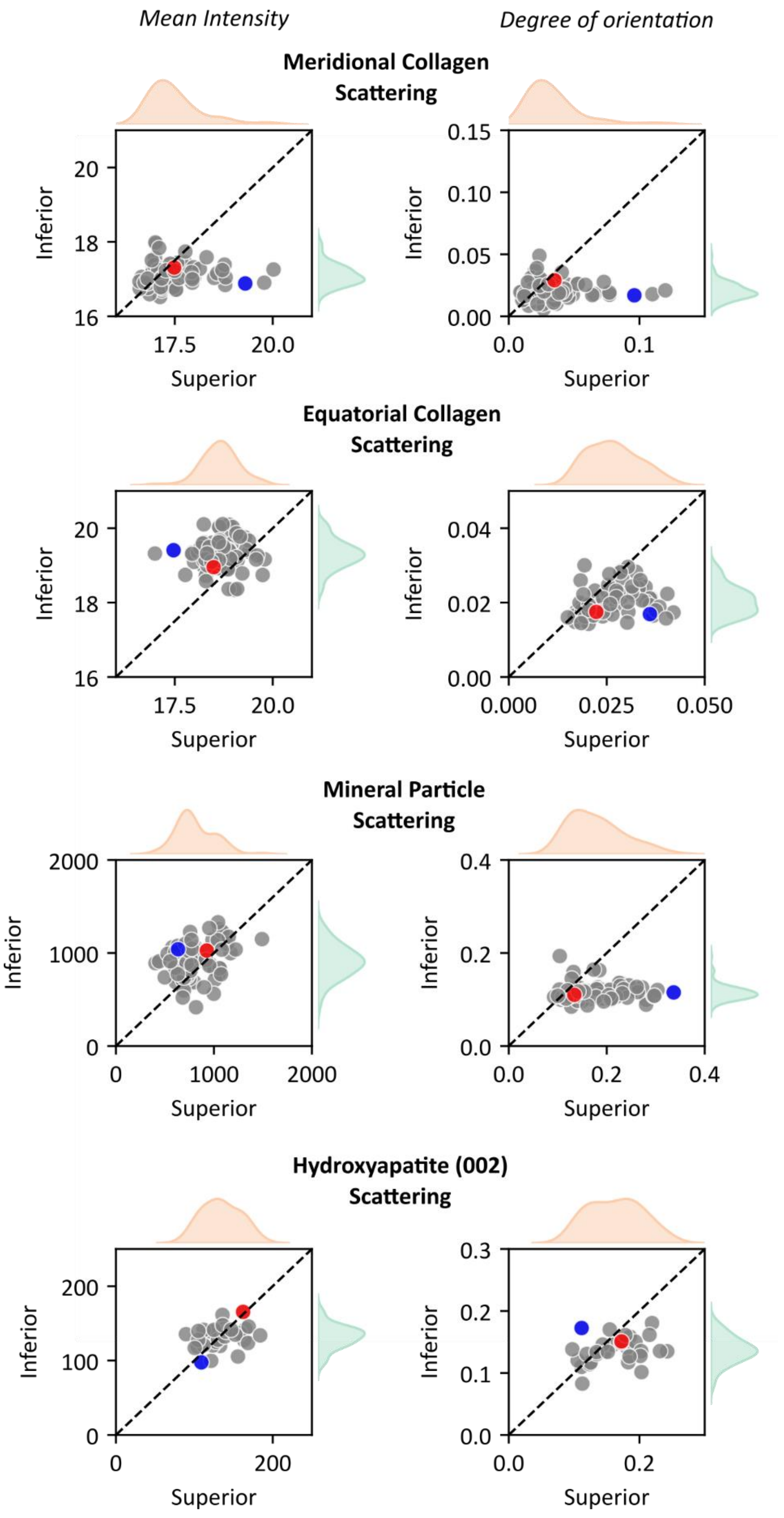


*Figure S10: Mean Intensity (left column) and 2D degree of orientation (right column) of the 4 analyzed scattering signals: Meridional collagen scattering (row1), equatorial collagen scattering (row 2), mineral particle scattering (row3), hydroxyapatite 002 scattering (row 4). For each femoral neck (n = 78), a scatter point represents the median value in the superior quadrant (x-axis) and inferior quadrant (y-axis). The blue and red points indicate the sample pairs from figure 4.*

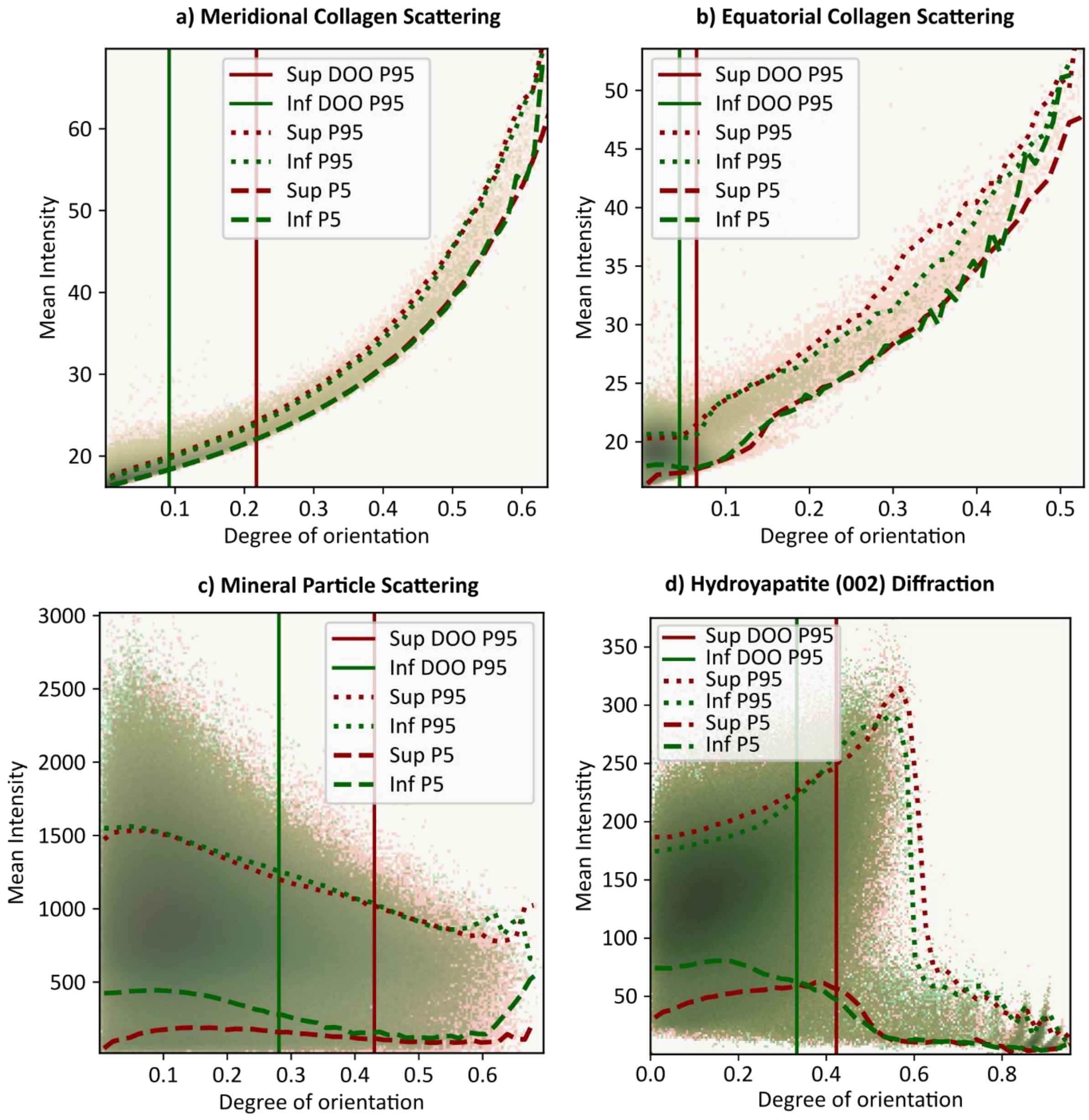


*Figure S11: Correlations between degree of orientation and mean intensity for the 4 analyzed scattering signals for the 2D dataset (4.3M pixels), separated by anatomical location. The 95$^{th}$ percentiles in the degree of orientation distributions (DOO) and the 5$^{th}$ and 95$^{th}$ percentiles in the intensity per degree of orientation bin are shown for each anatomical location. The meridional collagen scattering (a) shows the strongest correlation, suggesting it has a high dependency on the out-of-plane angle. The greater variation in the other signals (b-d) suggest greater structural variation in the respective scatterers. All 4 signals suggest a systematic change in out-of-plane angle between the two anatomical locations, considering the shift towards higher degree of orientations (Sup and Inf DOO p95 lines). The higher mean intensity in the mineral particle scattering (c) In plot (d), the data in the bottom right is caused by scratches in the samples and was not used in the statistical analysis.*

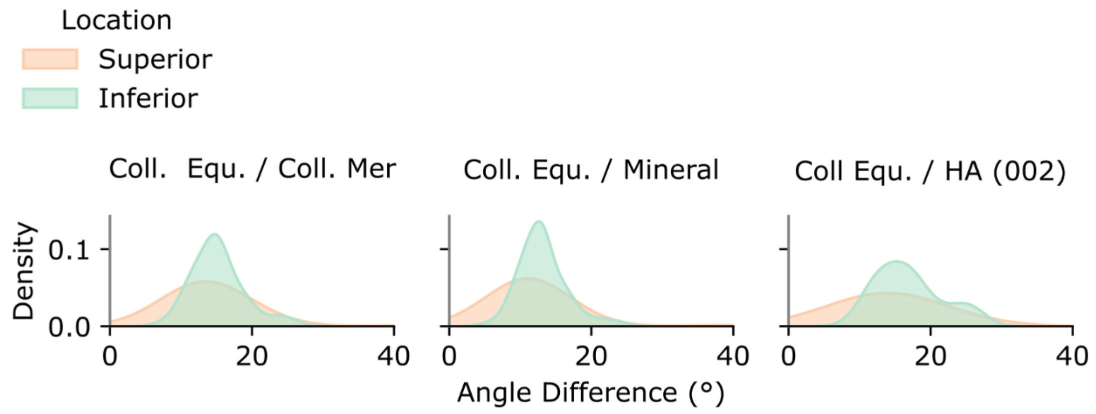


*Figure S12: Orientation misalignment of the collagen equatorial scattering with the collagen meridional scattering, mineral particle and hydroyapatite (002) scattering. The misalignment is greater on the inferior side in all cases, but the distributions are broader on the superior side.*

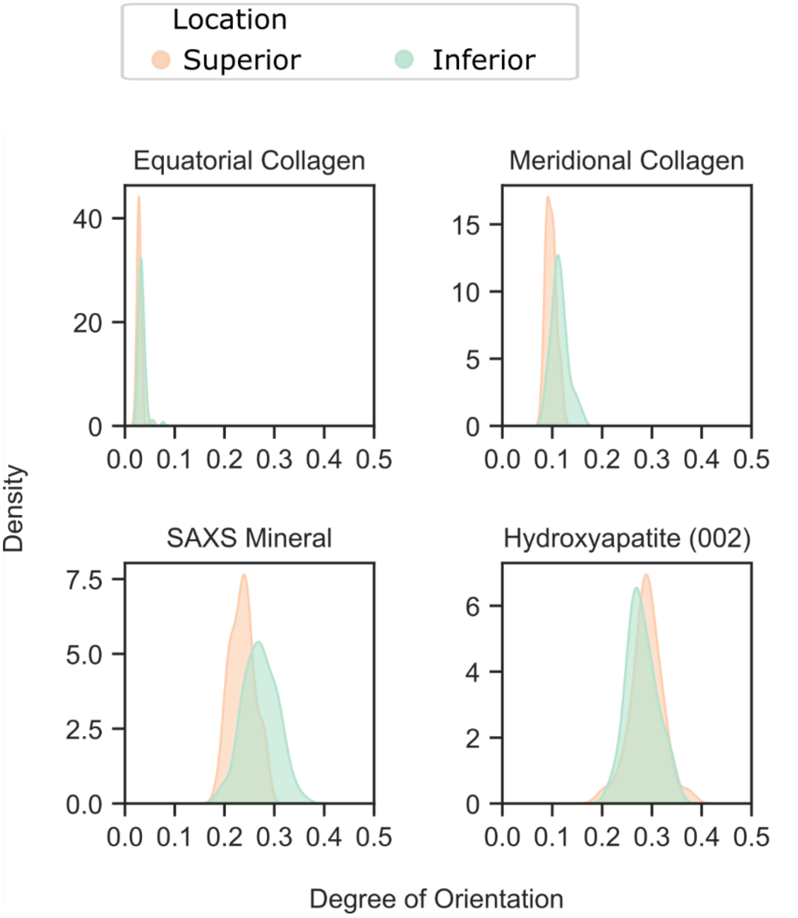


*Figure S13: 2D degree of orientation of the 4 analyzed scattering signals, per anatomical location. Both collagen signals display very low degree of orientation ($\lesssim 0.15$), due to the predominantly longitudinal MCF orientation. Both mineral signals show slightly higher degree of orientation ($\sim 0.2 - 0.4$), indicating a larger amount of oriented mineral platelets and HA crystals in the transversal plane. This could indicate the presence of extrafibrillar mineral being oriented due to the osteonal growth direction. The differences between anatomical locations reflect the out-of-plane angle difference.*

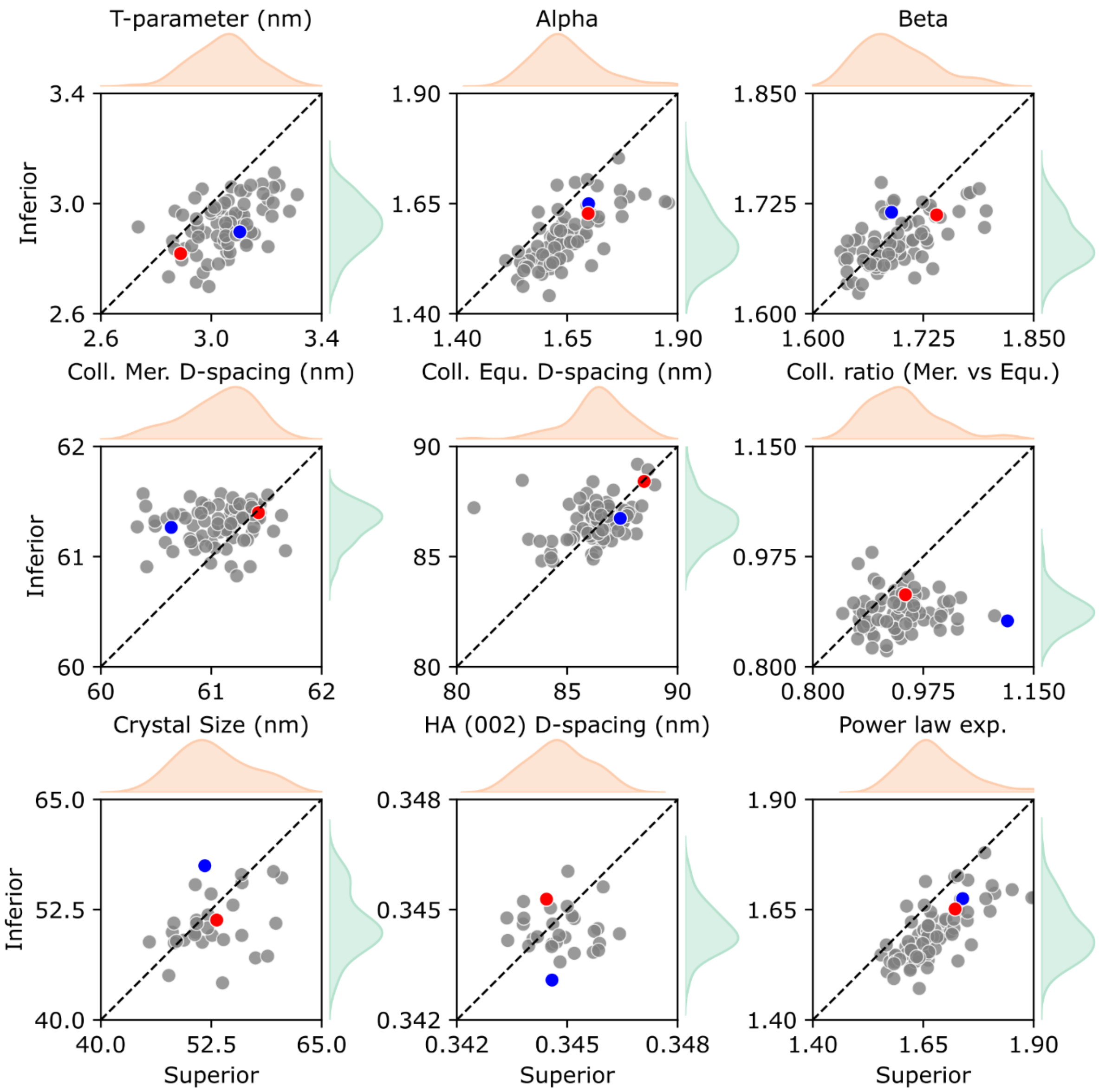


*Figure S14: Structural parameters measured in the 2D scanning SAXS/WAXS dataset. For each femoral neck (n = 78), a scatter point represents the median value in the superior quadrant (x-axis) and inferior quadrant (y-axis). The blue and red points indicate the sample pairs from figure 4.*

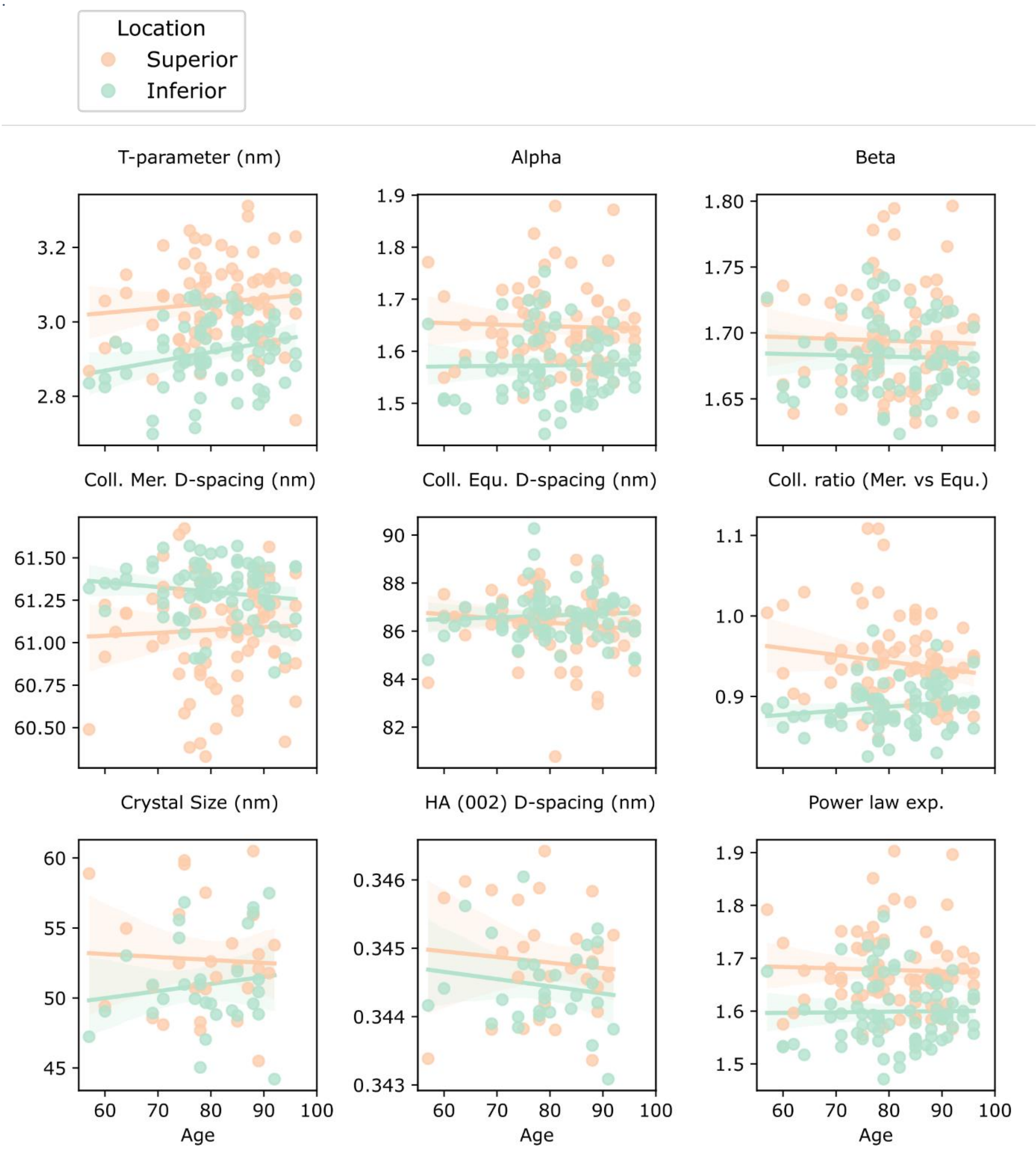


*Figure S15: Correlations between structural parameters and age of donors, separated by anatomical location. No significant correlations were noted, p-values are shown in Table S3.*

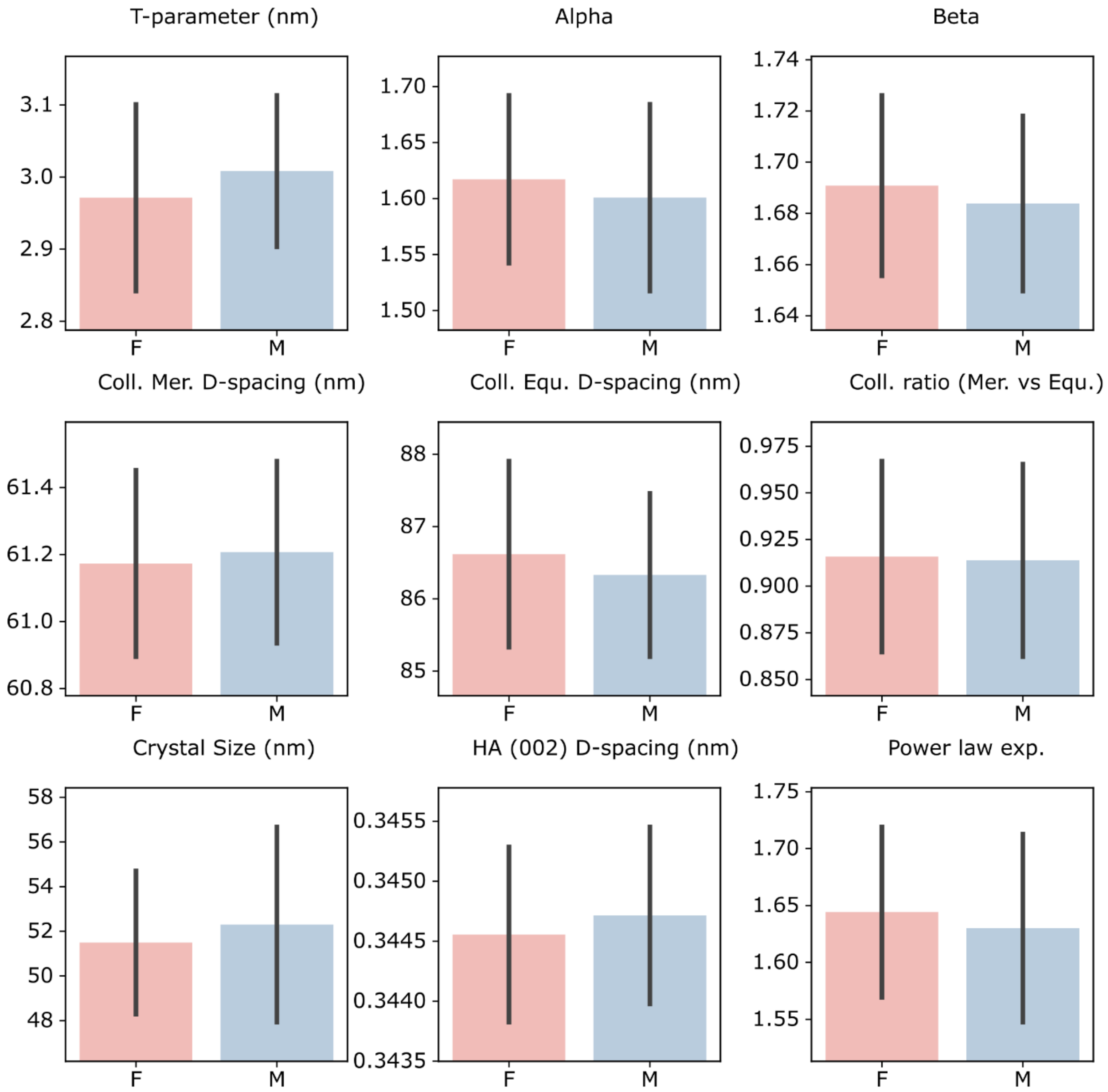


*Figure S16: Correlations between structural parameters and sex of the donors. No significant correlations were noted, p-values are shown in Table S3.*

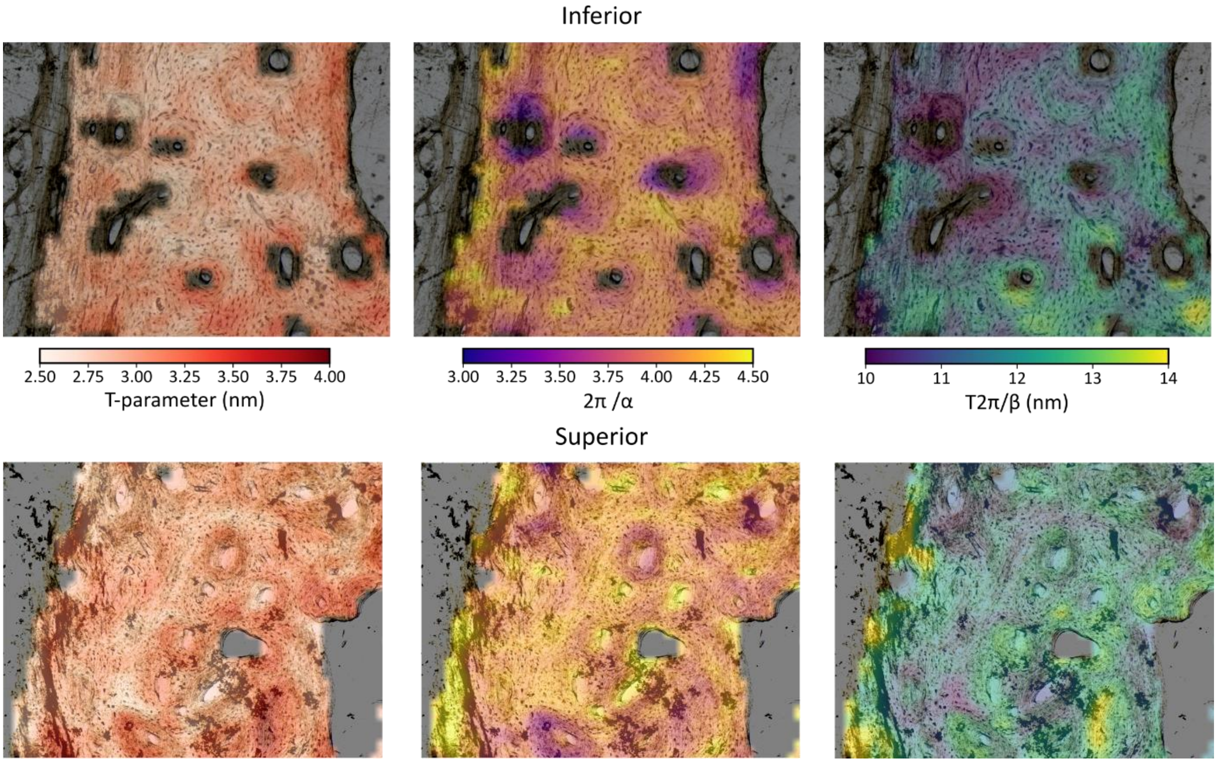


*Figure S17: Mineral thickness (T-parameter) and arrangement parameters (short-range ordering 2π/α and inter-platelet distance T2π/β) for an inferior and superior sample presenting different microstructural characteristics. The inferior side presents osteons with lower short-range order (lower 2π/α) and lower inter-platelet distance, while the osteons on the superior side display more variation.*
